# Human effector CD8^+^ T cells with an exhausted-like phenotype control tumor growth *in vivo* in a humanized tumor model

**DOI:** 10.1101/2023.10.11.561856

**Authors:** Juliane Mietz, Meike Kaulfuss, Lukas Egli, Lennart Opitz, Christian Münz, Obinna Chijioke

## Abstract

**Background:** Humanized tumor models could be particularly valuable for cancer immunotherapy research, as they may better reflect human-specific aspects of the interfaces between tumor and immune system of human cancer. However, endogenous antitumor immunity in humanized models is still largely undefined.

**Methods:** We established a novel autologous humanized mouse tumor model by using NSG mice reconstituted with human immune cells from hematopoietic progenitors and tumors generated from transformed autologous human B cells. We demonstrate growth of solid lymphoid tumors after subcutaneous implantation, infiltration by endogenous human immune cells and immunocompetence of the model.

**Findings:** We found human T cell subsets described in human cancer, including progenitor exhausted (T_pex_), terminally exhausted (T_ex-term_) and tissue-resident (T_RM_) cells in tumor-bearing humanized mice with accumulation of T_ex-term_ and T_RM_ in the tumor. In addition, we identified tumor-reactive CD8^+^ T cells through expression of CD137. This subpopulation of de novo arising human CD137^+^ CD8^+^ T cells displayed a highly proliferative, fully activated effector and exhausted-like phenotype with enhanced expression of activation and exhaustion markers like PD-1, CD39, CD160, TIM-3, TIGIT and TOX, the senescence marker CD57 (*B3GAT1*) and cytolytic effector molecules such as *PRF1*, *GZMH* and *NKG7*. Moreover, these CD137^+^ CD8^+^ T cells exhibited tumor-specific clonal expansion and presented signature overlap with tumor-reactive CD8^+^ T cells described in human cancer. We demonstrate superior anticancer activity of this exhausted-like human CD8^+^ T cell subset by adoptive transfer experiments using recipients bearing autologous human tumors. Mice adoptively transferred with CD137^+^ CD8^+^ T cells showed reduced tumor growth and higher CD8^+^ T cell tumor infiltration, correlating with control of human tumors.

**Interpretation:** We established an immunocompetent humanized tumor model, providing a tool for immunotherapy research and defined effective anticancer activity of human effector CD8^+^ T cells with an exhausted-like phenotype, supporting clinical exploration of such cells in adoptive T cell therapies.

**Funding:** Swiss Cancer Research foundation.

**Research in context:** *Evidence before this study:* Antitumor immune responses and outcome of immunotherapeutic interventions are not always consistent between mouse models of cancer and data available in humans. This may be due to species-specific differences, therefore models with a potential for better translatability are needed, such as humanized mouse models. However, there is limited data on human antitumor T cell immunity in humanized mice.

*Added value of this study:* In this study, we established an immunocompetent humanized tumor model that recapitulates hallmarks of human antitumor T cell responses, offering the possibility for further translational investigation of the interface between human tumors and endogenous anticancer immunity. Furthermore, using functional *in vitro* assays and adoptive transfer, our study demonstrates the key importance of human effector CD8^+^ T cells with an activated and exhausted-like phenotype in the antitumor immune response.

*Implications of all the available evidence:* The autologous humanized tumor model provided in this study can serve as a tool to elucidate human-specific immune features. By bridging a gap between syngeneic mouse tumor models and human-specific antitumor immune responses, the model may help open up avenues for greater translatability of preclinical data. Our findings suggest that exhausted-like effector CD8^+^ T cells can be harnessed for clinical development of adoptive T cell therapies.

## Introduction

The characterization of phenotypic traits that can faithfully identify tumor-specific CD8^+^ T cells with functional anticancer activity is an area of intense research. Recent studies have shown tumor-reactive CD8^+^ T cells to be rare in patients with various cancers and thus challenging to explore (1,2). Preclinical studies in mice have shown that CD8^+^ T cells displaying an exhausted-like phenotype have decreased effector function (3). Reinvigoration of exhausted antitumor T cells is thought to be a key component of response to immune checkpoint blockade. Nevertheless, studies in individuals with cancer have correlated exhausted-like phenotypic states of CD8^+^ T cells with tumor reactivity and clinical benefit (4,5).

Undoubtedly, tumor models using inbred and transgenic mice have contributed substantially to our understanding of tumor immunobiology and have been fundamental to the development of cancer immunotherapies that are now used on a routine basis in the clinic (6,7). However, human clinical studies (8) have shown that the outcome of cancer immunotherapies can significantly diverge from those of mouse models of cancer (9). This might be due to a relative lack of genetic diversity of mouse models compared to humans (10) but also to differences in aspects of gene expression networks (11), genomic organization (12) and tissue structure of the immune system of mice and humans (13). Xenograft mouse models using adoptive transfer of human immune cells rely on immunodeficient hosts, precluding analysis of the impact of endogenous adaptive antitumor immunity. Furthermore, in most preclinical studies investigating novel immunotherapy targets, monoclonal T cells with a single antigen specificity are explored (14,15) while polyclonal T cell responses might be a biomarker for favorable patient outcome to immunotherapy (16) and thus be more informative.

Complementing existing mouse tumor models, the use of mice with reconstituted human immune system components (HIS mice) holds hitherto underexplored translational potential to advance human cancer immunology (17). Yet, endogenous antitumor immunity in these models has remained largely undefined, mainly due to allogeneic reactivity of the reconstituted human T cells against allogeneic tumors (18), making a distinction of alloreactivity from tumor-specific immune responses challenging.

In this study, we demonstrate that reconstitution of human immune system components across independent donors can mount endogenous antitumor T cell responses to autologous human tumor challenge in HIS mice. A polyclonal population of human effector CD8^+^ T cells with an activated and exhausted-like phenotype that arises de novo in tumor-bearing HIS mice, resembling exhausted tumor-reactive lymphocytes found in patients with cancer, is able to control autologous human tumor growth *in vivo*. Thus, we provide a tool to investigate human antitumor T cell immunity and direct evidence of clinically relevant anticancer activity of expanded human tumor-reactive effector CD8^+^ T cells with an activated and exhausted-like, non-progenitor-like phenotype.

## Methods

### HIS & NSG mice

NSG (NOD.Cg-Prkdc<scid>Il2rg<tm1wjl>/SzJ (#005557)) mice were purchased from The Jackson Laboratory and bred and housed under specific pathogen-free conditions at the Laboratory Animal Services Center (LASC) Zurich, University of Zurich. To generate HIS mice, newborn pups (1-5 days old) were irradiated (1 Gy) and intrahepatically injected with 0.2 × 10^6^ human CD34^+^ hematopoietic progenitor cells derived from human fetal liver (Advanced Bioscience Resources, USA). CD34^+^ cells were isolated by MACS technology (Miltenyi, Cat. 130-046-703) following the manufacturer’s instructions. At 3 months of age, reconstitution of human immune cells was assessed in PBMC from tail vein blood by flow cytometry (19). Reconstitution analysis included staining for human CD45, CD3, CD4, CD8, CD19, NKp46 and HLA-DR. Only mice with a sufficiently high frequency of human CD45 (>25% of leukocytes) were included in the experiments. Both female and male mice were used in the experiments and no sex-based analysis was performed. All animal experiments were performed according to approved licenses by the veterinary office of the canton of Zurich, Switzerland (ZH049/20 and ZH067/2023).

### LCL generation and tumor model

LCL were generated from CD19^+^ B cells derived from the same autologous human fetal liver tissue from which the CD34^+^ cells for reconstitution of the human immune system in the same experiment were derived. CD19^+^ cells were isolated from the HFL tissue’s CD34^−^ fraction by MACS technology (Miltenyi, Cat. 130-050-301) according to the manufacturer’s instructions. 0.25 × 10^6^ CD19^+^ human B cells were transformed by infection with Epstein-Barr virus (EBV B95-8; produced as previously described (19)) at an MOI of 0.1 - 0.15 and cultured in RPMI/ 10% FCS/ 1% Penicillin/ Streptomycin.

LCL tumors were injected subcutaneously into the left flank under isoflurane narcosis. For generation and phenotyping of tumor-reactive T cells, 5 × 10^6^ autologous LCL tumor cells were resuspended in PBS and right before injection mixed in a 1:1 V/V ratio with Corning® Matrigel® Growth Factor Reduced (GFR) Basement Membrane Matrix (Milian, Cat. 354230).

For adoptive cell transfer experiments, 2 × 10^6^ LCL were injected s.c. in a 1:1 V/V mix with Matrigel and three days after tumor injection, 2 × 10^6^ T cells (HIS recipient mice) or 10 × 10^6^ (NSG recipient mice) were adoptively transferred by tail vein injection. Tumor size was monitored by calipering (3x/week or daily). Tumor volume was calculated as volume [mm^3^] = length [mm] × width^2^ [mm^2^] × 0.52. General health was monitored by weighing and health parameter scoring (3x/week or daily, according to animal license). PBMC composition and expansion of adoptively transferred T cells were monitored by weekly tail vein bleeding and flow cytometric analysis. White blood cell (WBC) counts were measured from blood with an automatic cell counting machine (DxH 500, Beckman Coulter). Animals were euthanized when they met pre-defined termination criteria defined in the animal license.

### T cell isolation and expansion

Spleens from LCL tumor-bearing and naïve HIS mice were harvested 16-18 days after s.c. tumor injection. Splenocytes were labelled with fluorescently labelled antibodies and sorted by fluorescent activated cell sorting (FACS). CD137^+^, CD137^−^ and CD137^−^PD-1^−^ CD8^+^ T cells were sorted from tumor-bearing HIS mice and bulk CD8^+^ T cells were isolated from naïve HIS mice. Sorted T cells were expanded by a rapid expansion protocol (REP), as described elsewhere (20). REP medium consisted of a 1:1 mix of RPMI (Gibco, Cat. 7001612) and X-Vivo (Lonza, Cat. BE02-060F) containing 10% FCS (Biochrome, Cat. S0615-500ML) and 1% Penicillin/ Streptomycin (Thermo Fisher, Cat. 7001592). After sorting, T cells were cultured in REP medium with 30 ng/µl OKT3 (Miltenyi, Cat. 130-093-387), 3000 IU/ml IL-2 (Peprotech, Cat. 200-02) and a 200-fold excess of irradiated (50 Gy) allogeneic PBMC derived from 3 healthy donors at a total cell density of 5 × 10^6^ cells/ml. Starting at day 7, cells were adjusted with REP medium to 0.5-1 × 10^6^ cells/ml and fresh IL-2 (3000 IU/ml) was added every second day. Cells were expanded for a total of 14-16 days until use for further experiments.

### Cell isolation from organs

For reconstitution analysis and weekly bleeding, 100-150 µl of tail vein blood was collected into EDTA tubes (BD Microtainer, BD, Cat. 365975) and erythrocytes were lysed by incubation with lysis buffer. Leukocytes were stained for flow cytometric analysis. For PBMC isolation on the day of sacrifice, blood was harvested by heart puncture and erythrocytes were lysed with erythrocyte lysis buffer. For splenocyte isolation, spleens were removed and meshed through a 70 µm cell strainer in PBS and lymphocytes were isolated by Ficoll-Paque (GE Healthcare, 17-5442-03). Single cell suspensions were washed and stained for flow cytometric analysis. For tumor-infiltrating lymphocyte (TIL) isolation, tumors were removed from the flank and skin was removed from the tumors. Tumors were cut into small pieces with scissors and then incubated with digestion mix (DMEM (Life technologies, Cat. 7001566) containing 2% FCS (Biochrome, Cat. S0615-500ML), 1 mg/ml Collagenase IV (Roche, Cat. 7002219), 10ug/ml DNase I (Roche, Cat. 7002221) and 1.2 mM CaCl2; approx. 1.5 ml/tumor, incubation for 45 min, 37°C). The tumor digest was diluted with 20 ml RPMI or PBS and meshed over a 70 µm cell strainer. Single cell suspensions were washed and then stained for flow cytometric analysis.

### Flow cytometry

For surface staining, cells were washed in PBS and stained with an antibody master mix for 20 min at 4°C in the dark. For intracellular staining, cells were first stained for surface antigens and fixable live/dead marker, then fixed and permeabilized using the BD Pharmingen™ Transcription Factor Buffer Set (BD, Cat. 562574) according to the manufacturer’s instructions, then stained for intracellular or intranuclear antigens for 50-60 minutes. Cells were washed in PBS and acquired on a BD LSRFortessa™ or Cytek Aurora.

Antibodies for reconstitution analysis of HIS mice (all antibodies are anti-human): CD45-Pacific Blue (Biolegend, HI30, Cat. 304029), CD3-PE (Biolegend, UCHT1, Cat. 300408), CD4-APC-Cy7 (Biolegend, RPA-T4, Cat. 300518), CD8-PerCP (Biolegend, SK1, Cat. 344708), HLA-DR-FITC (Biolegend, L243, Cat. 307604), CD19-PE-Cy7 (Biolegend, HIB19, Cat. 302216), NKp46-APC (BD, 9-E2, Cat. 558051), Zombie Aqua™ Fixable Viability Kit (Biolegend, Cat. 423101).

Antibodies for flow cytometry-based sorting of T cells: CD3-FITC (Biolegend, UCHT1, Cat. 300406), CD4-APC-Cy7 (Biolegend, RPA-T4, Cat. 300518), CD8-BV650 (Biolegend, SK1, Cat. 344730), CD45-Pacific Blue (Biolegend, HI30, Cat. 304029), CD137-APC (Miltenyi, REA765, Cat. 130-110-901), PD-1-PE-Dazzle (Biolegend, EH12.2H7, Cat. 329940), Zombie Aqua™ Fixable Viability Kit (Biolegend, Cat. 423101).

Antibodies for phenotyping in Panel 1: CD45-Pacific Blue (Biolegend, HI30, Cat. 304029), CD3-PerCP-Cy5.5 (Biolegend, UCHT1, Cat. 300429), CD4-APC-Cy7 (Biolegend, RPA-T4, Cat. 300518), CD8-BV650 (Biolegend, SK1, Cat. 344730), CD45RA-BV785 (Biolegend, HI100, 304139), CD62L-FITC (BD, DREG-56, Cat. 555543), CD19-PE-Cy7 (Biolegend, HIB19, Cat. 302216), HLA-DR-BV605 (BD, g46-6, Cat. 562845), CD39-BV711 (Biolegend, A1, Cat. 328228), CD69-Alexa Fluor 700 (Biolegend, FN50, Cat. 310921), PD-1-PE-Dazzle (Biolegend, EH12.2H7, Cat. 329940), TCF1-PE (Biolegend, 7F11A10, Cat. 655207), CD137-APC (Miltenyi, REA765, Cat. 130-110-901), Zombie Aqua™ Fixable Viability Kit (Biolegend, Cat. 423101); in Panel 2: CD3-FITC (Biolegend, UCHT1, Cat. 300406), CD4-PE-Cy7 (Biolegend, RPA-T4, Cat. 300512), CD8-BV650 (Biolegend, SK1, Cat. 344730), CD45-BV711 (Biolegend, HI30, Cat. 304050), CD137-APC-Fire750 (Biolegend, 4B4-1, Cat. 309834), Granzyme B-APC-Alexa700 (Biolegend, QA16A02, Cat. 372222), Ki-67-BV605 (Biolegend, Ki-67, Cat. 350522), PD-1-PE-Dazzle (Biolegend, EH12.2H7, Cat. 329940), Perforin – BV421 (Biolegend, dG9, Cat. 308122), TCF1-PE (Biolegend, 7F11A10, Cat. 655207), TOX-APC (Miltenyi, REA473, Cat. 130-118-335), Zombie Aqua™ Fixable Viability Kit (Biolegend, Cat. 423101); in Panel 3: CD45-BUV395 (BD, HI30, Cat. 563792), CD3-Pacific Blue (Invitrogen, S4.1, Cat. MHCD0328), CD4-BUV495 (BD, SK3, Cat. 612936), CD8-BUV563 (BD, RPA-T8, Cat. 612914), CD19-PE-Cy5 (Biolegend, HIB19, Cat. 302210), CD45RA-BV785 (Biolegend, HI100, 304139), CD62L-BV605 (Biolegend, DREG-56, Cat. 304834), CD39-PE-Fire810 (Biolegend, A1, Cat. 328245), CD69-BUV805 (BD, FN50, Cat. 748763), CD137-APC (Miltenyi, REA765, Cat. 130-110-901), PD-1-PE-Dazzle (Biolegend, EH12.2H7, Cat. 329940), HLA-DR-FITC (Biolegend, LN3, Cat. 327006), TCF1-PE (Biolegend, 7F11A10, Cat. 655207), CD160-PerCP-Cy5.5 (Biolegend, BY55, Cat. 341210), LAG3-BV650 (Biolegend, 11C3C65, Cat. 369316), TIM3 – Alexa Fluor 700 (R&D Systems, Clone 344823, Cat. FAB2365N), Galectin-9-PE-Cy7 (Biolegend, 9M1-3, Cat. 348916), TIGIT-APC-Cy7 (Biolegend, A15153G, Cat. 372734), Ki-67-BV510 (Biolegend, Ki-67, Cat. 350518), CD103-BV711 (Biolegend, Ber-ACT8, Cat. 350222), LIVE/DEAD^TM^ Fixable Blue Dead Cell Stain Kit (Invitrogen, Cat. L23105).

### Co-expression analysis

For co-expression analysis of flow cytometric data, cells were manually gated on live, singlet human CD8^+^ T cells. Cells were analyzed using CRUSTY webtool (21), where clustering was performed using Phenograph algorithm and default settings for UMAP generation.

### IFN-γ ELISpot and TNFα ELISA

For IFN-γ ELISpot, ELISpot plates (Merck, Cat.MAIPN4550) were activated with 35% ethanol, washed and coated overnight with anti-IFN-γ coating antibody (mAb-1-D1K, Mabtech, Cat. 3420-3-1000). Plates were washed and T cells were co-cultured with autologous LCL in RPMI/ 10% FCS/ 1% Pen/Strep at an effector: target ratio of 5:1 for 16-24 hours. After incubation, supernatant was harvested, stored at −20°C and used for TNFα ELISA. Wells were washed and incubated for 2 hours with biotinylated anti-IFN-γ detection antibody (mAb-7-B6-1-Biotin, Mabtech, Cat. 3420-6-250). Afterwards, wells were washed and incubated with Streptavidin-ALP (Mabtech, Cat. 3310-10) for 1 hour. Wells were then washed again and spots were developed with filtered substrate solution BCIP/ NBT-plus (Mabtech, Cat. 3650-10). When distinct spots were visible, the plate was washed and dried overnight. Plates were acquired with an ELISpot reader (AID classic, AID GmbH) and spots were automatically counted using AID ELISpot 8.0 software (AID GmbH).

TNFα ELISA was performed with the supernatant from T cell and LCL co-culture as described above using the human TNFα ELISA^BASIC^ kit (Mabtech, Cat. 3512-1H-6), following manufacturer’s instructions. Plates were acquired using an ELISA reader (infinite M200Pro, Tecan) and analyzed with the i-control 2.0 software (Tecan).

### Immunohistochemistry

Tumors were fixed in 4% paraformaldehyde and embedded in paraffin. Sample embedding, preparation and staining was performed by the Pathology department of the University Hospital Basel. Slides were acquired on an automated slide scanning brightfield microscope (Vectra 3, Akoya Biosciences) and quantified using InForm software (Akoya Biosciences).

### Bulk RNAseq and TCR profiling

RNA was extracted from 5 × 10^6^ T cells using the RNeasy® Mini Kit (Qiagen, Cat. 74106). RNA extraction was performed following the instructions from the “Purification of Total RNA from Animal Cells Using Spin Technology” protocol given in the RNeasy® Mini Handbook, including homogenization with QIAshredders (Qiagen, Cat. 79654) and on-column DNA digestion with the RNase-free DNase Set (Qiagen, Cat. 79254). All further steps were performed by the Functional Genomics Center Zurich (FGCZ). The quality of the isolated RNA was determined with a Qubit® (1.0) Fluorometer (Life Technologies, California, USA) and a Bioanalyzer 2100 (Agilent, Waldbronn, Germany). Only those samples with a 260 nm/280 nm ratio between 1.8–2.1 and a 28S/18S ratio within 1.5–2 were further processed. The TruSeq RNA Sample Prep Kit v2 (Illumina, Inc, California, USA) was used in the succeeding steps. Briefly, total RNA samples (600 ng) were poly A enriched and then reverse-transcribed into double-stranded cDNA. The cDNA samples were fragmented, end-repaired and polyadenylated before ligation of TruSeq adapters containing the index for multiplexing fragments containing TruSeq adapters on both ends were selectively enriched with PCR. The quality and quantity of the enriched libraries were validated using Qubit® (1.0) Fluorometer and the Caliper GX LabChip® GX (Caliper Life Sciences, Inc., USA). The product is a smear with an average fragment size of approximately 260 bp. The libraries were normalized to 10nM in Tris-Cl 10 mM, pH8.5 with 0.1% Tween 20. After library quantification, libraries were prepared for loading accordingly to the NovaSeq workflow with the NovaSeq6000 Reagent Kit (Illumina, Catalog No. 20012865). Cluster generation and sequencing were performed on a NovaSeq6000 System with a run configuration of single end 100bp. RNA-seq data analysis consisted of the following steps: The raw reads were first cleaned by removing adapter sequences, trimming low quality ends, and filtering reads with low quality (phred quality <20) using Fastp (Version 0.20) (22). Sequence pseudo alignment of the resulting high-quality reads to the human reference genome (build GRCh38.p13 and GENCODE gene models release 37) and quantification of gene level expression was carried out using Kallisto (Version 0.46.1) (23). Differential expression was computed using the generalized linear model implemented in the Bioconductor package DESeq2 (R version: 4.2.2, DESeq2 version: 1.38.1) (24). Genes showing altered expression with adjusted (Benjamini and Hochberg method) p-value < 0.05 were considered differentially expressed. RNA-seq expression data underwent preprocessing to remove duplicated genes. Subsequently, we utilized the CreateSeuratObject() function within the Bioconductor package Seurat (R version: 4.2.2, Seurat version: 5.0.1) (25) to construct a Seurat object, facilitating downstream analyses. Normalization of the expression data was performed using the NormalizeData() function, with experimental group assignments explicitly indicated within the Seurat object. To delineate the gene signatures characterizing distinct T cell phenotypes, we curated pertinent signatures from previous studies. Genes absent in our expression data were systematically filtered out. Next, employing the AddModuleScore() function, we computed module scores for each gene signature across experimental groups and values were scaled for heatmap visualization. For the TCR analysis the read alignment to the Human genome was performed with STAR (v2.7.10b) (26). TCR sequences (CDR3 of TRA, TRB, TRG and TRD) were reconstructed from bulk RNAseq data using TRUST4 (v1.0.10) (27). Out-of-frame sequences were excluded from the analysis. TCR repertoire analysis was performed using the R package Immunarch (28).

### Statistics

Statistical analysis was performed using Prism 10 Software (GraphPad Software, Inc). Figure legends indicate which statistical test was performed.

### Study approval

Animal experiments were conducted according to licenses approved by the veterinary office of the canton of Zurich, Switzerland (ZH049/20 and ZH067/2023).

### BioRender

Schematics were generated with a licensed version of BioRender.

## Author contributions

Conceptualization, J.M. and O.C.; Methodology, J.M., M.K. and O.C.; Investigation, J.M., M.K., L.E., L.O. and O.C.; Writing – Original Draft:, J.M. and O.C.; Writing – Review & Editing, J.M. C.M. and O.C.; Funding Acquisition, O.C.; Resources, L.O. and C.M.; Supervision, O.C.

## Acknowledgments

We would like to thank Fernando Canale for help with gene signature analysis and Daniel E. Speiser for critically reading the manuscript. Funding: This research was supported by Swiss Cancer Research foundation (KFS-5292-02-2021 to O.C.).

## Data Sharing

Supplemental data are provided with this paper. All data are also available upon request.

## Results

### Establishment of an autologous HIS mouse tumor model

We established and characterized a novel tumor model in which immunocompetent, humanized mice are subcutaneously implanted with autologous human tumor cells. For this aim, we isolated CD34^+^ hematopoietic progenitor cells (HPC) and CD19^+^ B cells from the same human HPC donor tissue. CD34^+^ HPCs were engrafted into NSG mice to reconstitute components of the human immune system (from here on referred to as human immune system (HIS) mice), while CD19^+^ B cells were transformed into lymphoblastoid cell lines (LCL) by infection with the oncogenic Epstein-Barr virus (EBV) (**Figure 1A**).

**Figure 1:**
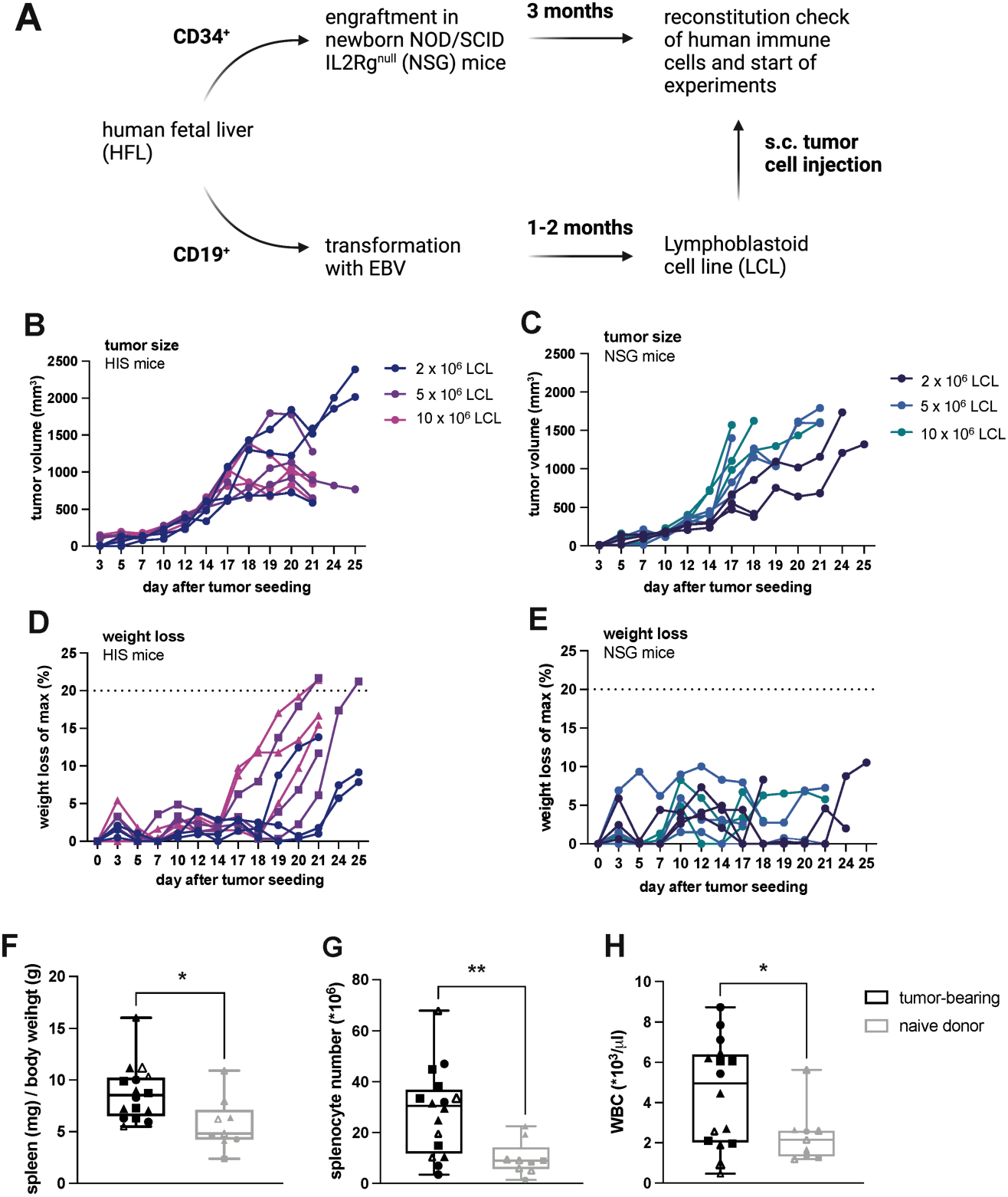
LCL tumor growth kinetics and weight loss in autologous HIS mice and NSG mice. LCL tumors were injected subcutaneously into the flank of mice and tumor size was assessed by calipering. **A**, schematic of generation of HIS mice and autologous tumor cells and subsequent injection in HIS mice. **B**, tumor volume in HIS mice after implantation of the indicated number of LCLs. **C,** Tumor volume in NSG mice after implantation of the indicated number of LCLs. (D, E) Relative weight loss calculated based on the maximum weight during the experiment after tumor cell implantation of the indicated number of LCLs in HIS (**D**) or NSG (**E**) mice. **F**, ratio of spleen weight (mg) to body weight (g), **G**, total splenocyte number and **H**, white blood cell count in HIS mice injected with 5 × 10^6^ autologous LCL tumor cells or non-tumor bearing HIS mice (naïve) sacrificed 15-20 days after tumor implantation. B-E, n=3 per tumor cell number, F-H, n(tumor-bearing)=16, n(naïve)=9 from 4 individual experiments, marked by individual symbols. Unpaired t-test, data shown as box and whiskers with the box from the 25-75 percentile and the median is shown as a line within the box; whiskers are shown from minimum to maximum data point.

To characterize LCL tumorigenicity and tumor growth in the presence or absence of an autologous human immune system, HIS mice reconstituted with HPCs autologous to the tumor or NSG mice were injected subcutaneously with LCLs. In both recipient strains, palpable tumors formed at the site of implantation. In HIS mice, in the presence of an endogenous immune system, tumors showed progressive growth until approximately day 18 and subsequently, a majority of tumors showed regression (**Figure 1B**). In contrast, in NSG mice without an endogenous immune system, tumors increased in size until termination criteria were reached (**Figure 1C**). These data suggested a functional antitumor response mediated by components of the reconstituted human immune system, i.e. immunocompetence, in our autologous HIS mouse tumor model.

### Endogenous antitumor immune response in HIS mice

To investigate the tumor regression that we observed in the presence of an autologous human immune system, we next characterized the immune response in tumor-bearing HIS mice. Around the same time of tumor regression, we found weight loss in HIS mice (**Figure 1D**), but not in tumor-bearing NSG mice (**Figure 1E**). This indicated immunopathology related to the endogenous human antitumor immune response, reminiscent of cancer cachexia (29) rather than an effect of tumor growth per se. Additionally, we found increases in spleen-to-body weight ratio, total splenocyte count and white blood cell count in tumor-bearing HIS mice (**Figure 1, F-H**), suggesting immune cell proliferation occurring in response to tumor growth. Furthermore, tumors in HIS mice showed infiltration by endogenous human immune cells, including CD8^+^ T cells (Supplemental Figure 1).

The frequency of human CD3^+^ T cells in tumor-bearing HIS mice in spleen (**Figure 2A**) and circulation (Supplemental Figure 2A) increased significantly compared to naïve HIS mice. Moreover, tumor-bearing HIS mice showed an increased frequency of CD8^+^ T cells, suggesting preferential expansion (**Figure 2A** and Supplemental Figure 2A). These CD8^+^ T cells exhibited differentiation towards an effector memory (T_EM_) phenotype and, to a lower degree, central memory (T_CM_) phenotype, whereas non-tumor bearing HIS mice had higher frequencies of naïve T cells (T_naïve_) (**Figure 2B** and Supplemental Figure 2B). Consistent with an exhausted effector-like phenotype, CD8^+^ T cells from tumor-bearing HIS mice also showed enhanced expression of the exhaustion/activation markers PD-1, CD39 and HLA-DR (**Figure 2C** and Supplemental Figure 2C) as well as trending towards increased TOX expression (**Figure 2D** and Supplemental Figure 2D), while exhibiting significantly reduced levels of the progenitor/stemness-related marker TCF1 (30) (**Figure 2D** and Supplemental Figure 2D).

**Figure 2:**
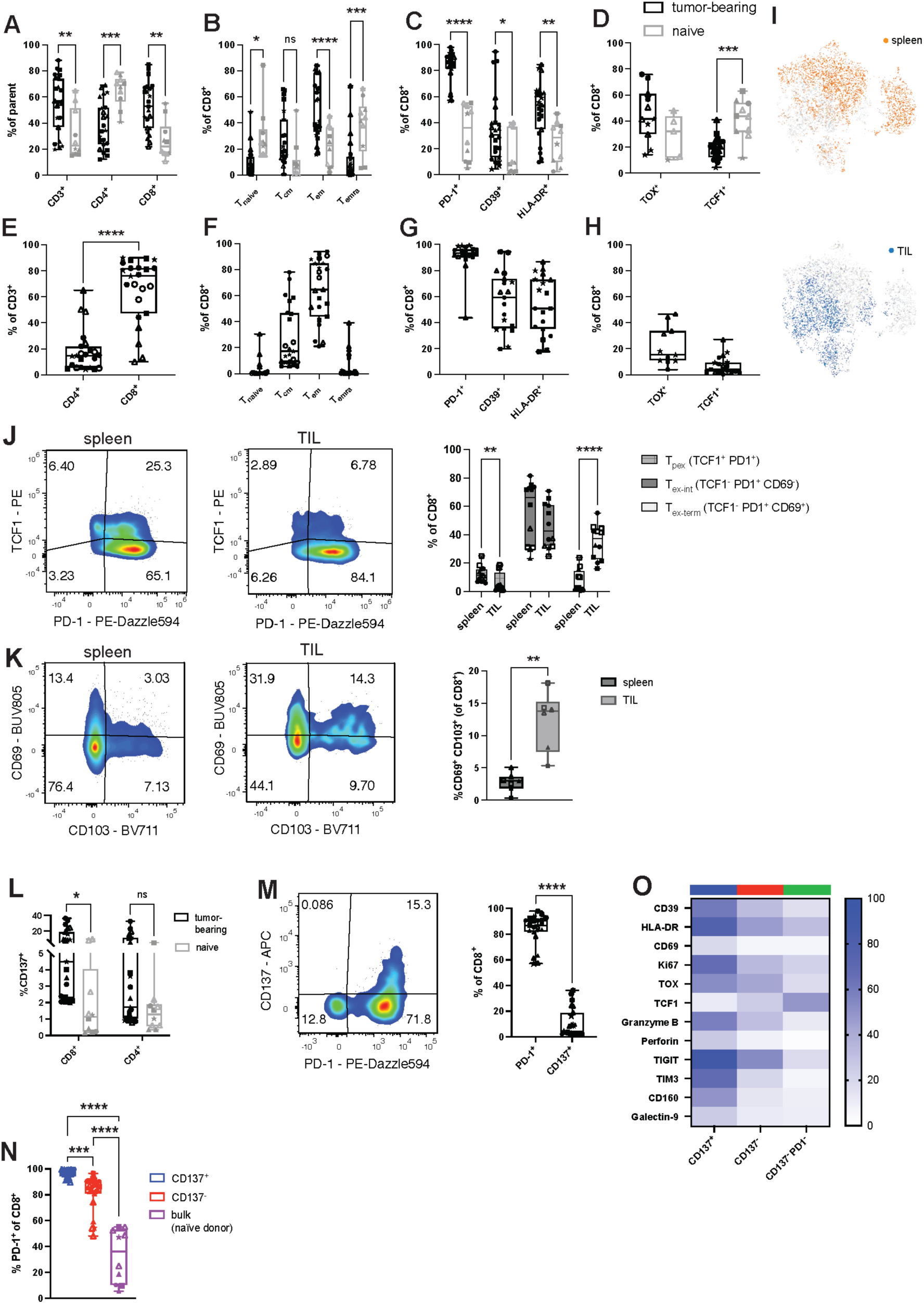
T cell response in tumor-bearing HIS mice. Splenocytes (A-D) and tumor-infiltrating lymphocytes (TIL) (E-H) were isolated from HIS mice 16-18 days after tumor implantation and analyzed by flow cytometry. **A,** frequency of splenic T cells in tumor-bearing or naïve HIS mice; parent population refers to frequency (%) of CD3^+^ T cells within human CD45^+^ cells and CD4^+^ and CD8^+^ T cells within CD3^+^ T cells. **B**, CD8^+^ T cell differentiation defined as T_naïve_ (CD45RA^+^CD62L^+^), T_CM_ (CD45RA^−^CD62L^+^), T_EM_ (CD45RA^−^CD62L^−^), T_EMRA_ (CD45RA^+^CD62L^−^) in tumor-bearing or naïve HIS mice. **C-D**, expression of indicated markers on CD8^+^ T cells from spleen of tumor-bearing or naïve HIS mice. **E,** frequency of CD4^+^ and CD8^+^ T cells within TILs, gated on total CD3^+^ T cells. **F,** CD8^+^ T cell differentiation within TILs. **G-H,** expression of indicated markers on CD8^+^ T cells within TILs. **I**, UMAP of CD8^+^ T cells from spleen and TIL of tumor-bearing HIS mice, colored for tissue of origin. **J**, expression of PD-1 and TCF1 in spleen and TIL of tumor-bearing HIS mice and quantification of CD8^+^ T_pex_ (TCF1^+^PD1^+^), T_ex-int_ (TCF1^−^PD1^+^CD69^−^), T_ex-term_ (TCF1^−^PD1^+^CD69^+^) in both tissues. **K**, expression of CD69 and CD103 in spleen and tumor of tumor-bearing HIS mice and quantification of T_RM_-like (CD69^+^CD103^+^) CD8^+^ T cells in both tissues. **L,** expression of CD137 on splenic CD8^+^ and CD4^+^ T cells of tumor-bearing or naïve HIS mice. **M,** expression and quantification of PD-1 and CD137 on splenic CD8^+^ T cells of tumor-bearing HIS mice. **N**, expression of PD-1 on splenic CD137^+^ or CD137^−^ CD8^+^ T cells in tumor-bearing HIS mice or bulk CD8^+^ T cells from naïve HIS mice. **O**, expression of individual markers on splenic CD8^+^ T cell populations from tumor-bearing HIS mice based on expression of CD137 and PD-1. Data are pooled from at least 3 independent experiments, n=10-23 mice per group. For each experiment, a different HPC donor was used for HIS mouse reconstitution and generation of autologous tumor. Significance by paired t-test, one-way ANOVA or 2way ANOVA, as appropriate. Data from individual experiments are indicated by different symbols, with individual mice from the same experiment indicated by the same symbol.

Similar to the increased CD8^+^ T cell frequencies in secondary lymphoid tissue (spleen) and circulation, tumors in HIS mice exhibited predominant infiltration by CD8^+^ T cells (**Figure 2E** and Supplemental Figure 1C). The majority of autologous tumor-infiltrating human CD8^+^ T cells had a T_EM_-like phenotype, with a lower proportion of T_CM_ and greatly reduced proportions of T_EMRA_ and naïve T cells (**Figure 2F**). Tumor-infiltrating CD8^+^ T cells displayed high expression of exhaustion and activation markers PD-1 (91.0%±12.1), CD39 (56.4%±22.8) and HLA-DR (53.9%±22.4) (**Figure 2G**), low levels of TCF1 (5.6%±5.0) (**Figure 2H**), while a fraction of tumor-infiltrating CD8^+^ T cells expressed TOX (21.7%±14.81) (**Figure 2H**). Dimensionality reduction by UMAP revealed obvious differences of CD8^+^ T cell landscapes between tumor-infiltrating lymphocytes (TIL) and spleen (**Figure 2I**). Together, the phenotype of human CD8^+^ T cells in tumor-bearing HIS mice was reminiscent of CD8^+^ T cell phenotypes described in human cancer (31,32).

Progenitor exhausted T cells (T_pex_; TCF1^+^PD-1^+^) play an important role for immunotherapy of human cancer. T_pex_ have been described as precursors of exhausted cytotoxic T cells (T_ex_; TCF1^−^PD-1^+,^, further subdivided into TCF1^−^PD-1^+^CD69^−^ intermediate exhausted (T_ex-int_) and TCF1^−^PD-1^+^CD69^+^ terminally exhausted (T_ex-term_) T cells (33)), which exhibit effector function but reduced self-renewal and long-term survival, making T_pex_ cells an important reservoir of antitumor T cells (33–35). Interestingly, we detected an increased fraction of CD8^+^ T_pex_ cells in secondary lymphoid tissue (spleen) of tumor-bearing HIS mice compared with tumor (**Figure 2J**). This distribution recapitulates human cancers, where a higher frequency of T_pex_ is found in lymph nodes compared to the tumor site (34). In addition, we found a significantly higher proportion of terminally exhausted CD8^+^ T cells (T_ex-term_; TCF1^−^PD-1^+^CD69^+^) within the tumor compared to the spleen, a secondary lymphoid organ (**Figure 2J**), again consistent with what has been described in human cancer (34,35).

Tissue-resident-memory (T_RM_) T cells (CD69^+^CD103^+^) within tumors are another T cell subset of interest for immunotherapy research. The presence of T_RM_ cells has been associated with a favorable outcome in certain cancer types and based on their expression of immune checkpoint molecules, tumor-reactive T_RM_ cells also constitute a target for immune checkpoint blockade (36). We detected T_RM_-like (CD69^+^CD103^+^) CD8^+^ T cells in tumor-bearing HIS mice and found a significantly higher abundance in tumor compared with spleen (**Figure 2K**).

The presence and identification of these human CD8^+^ T cell subsets, which are considered relevant in human cancer and are thought to play critical roles in response to immunotherapy, underscores the translational value of our humanized mouse model for immunotherapy research.

### Activated and exhausted-like, cytotoxic effector-like phenotype of human CD137^+^ CD8^+^ T cells

In human cancers, CD137 has been described as a marker for tumor-reactive T cells (37–39). We similarly identified a population of human CD137^+^ T cells in tumor-bearing mice (**Figure 2L**) and characterized the CD8^+^ T cell population further. Likewise, PD-1 has been reported to be a marker of tumor-reactive T cells in human cancers, both in circulation and within tumor-infiltrating lymphocytes (20,40,41). We observed that of splenic CD8^+^ T cells from tumor-bearing HIS mice, only a small proportion (10.9%±11.2) were CD137 positive, whereas the majority of CD8^+^ T cells (83.5%±12.1) expressed PD-1 on their surface (**Figure 2M**). Though there was a small but still significant difference between the overall high level of PD-1 expression on CD137^+^ CD8^+^ and CD137^−^ CD8^+^ T cells from tumor-bearing HIS mice, both these populations expressed PD-1 significantly more than bulk CD8^+^ T cells from naïve HIS mice (**Figure 2N**). These data suggested that CD137 alone might not be sufficient to delineate tumor-reactive T cell subsets. We therefore included a CD8^+^ T cell subset negative for both CD137 and PD-1 in our analysis to further explore and distinguish bona fide tumor-reactive from bystander CD8^+^ T cells in tumor-bearing HIS mice.

Indeed, compared to double negative CD137^−^PD-1^−^ CD8^+^ T cells, CD137^+^ CD8^+^ T cells and CD137^−^ CD8^+^ T cells displayed a T_EM_-like phenotype, while the double negative CD137^−^PD-1^−^population was enriched in naïve-like T cells (Supplemental Figure 2E). Within the splenic CD137^+^ CD8^+^ T cell population, we found the highest expression of the exhaustion marker CD39, which has been described as a marker of CD8^+^ T cells with enhanced tumor reactivity across different human cancers (1,42–46), the activation marker HLA-DR as well as the activation and residency-related marker CD69 (47), while this cell subset also had the strongest expression levels of Ki-67, indicating high proliferation (**Figure 2O** and Supplemental Figure 2F). Additionally, within the CD137^+^ CD8^+^ T cell population, we observed significantly increased expression of TOX, a transcription factor associated with exhaustion (48) and drastically reduced levels of the progenitor/stemness-related transcriptional regulator TCF1 (30) (**Figure 2O** and Supplemental Figure 2G). Several makers related to T cell exhaustion and activation (49,50) (TIGIT, TIM3, CD160, Galectin-9) were similarly upregulated on CD137^+^ CD8^+^ T cells (**Figure 2O** and Supplemental Figure 2H). Finally, CD137^+^ CD8^+^ T cells displayed the highest levels of the cytolytic molecules granzyme B and perforin compared with CD137^−^ CD8^+^ T cells or CD137^−^PD-1^−^ CD8^+^ T cells from tumor-bearing HIS mice (**Figure 2O** and Supplemental Figure 2I).

The CD137^+^ CD8^+^ T cell population contained a significantly higher proportion of T_ex-term_ cells in both spleen and tumor and a significantly lower frequency of T_pex_ in the tumor relative to the CD137^−^ CD8^+^ T cell population (Supplemental Figure 2J and 2K). As CD69 is expressed on subsets of both exhausted as well as T_RM_-like T cells (33), we quantified CD8^+^ T_RM_-like (CD69^+^CD103^+^) T cells and found no difference between CD137^+^ and CD137^−^ T cells in the spleen (Supplemental Figure 2L). In the tumor, where an overall higher proportion of T_RM_-like T cells was present compared to spleen (**Figure 2K**), the CD137^+^ CD8^+^ T cell population contained significantly fewer T_RM_-like T cells compared to CD137^−^ CD8^+^ T cells (Supplemental Figure 2M). Thus, the majority of CD137^+^ CD8^+^ T cells did not exhibit a T_RM_-like phenotype, but consisted of more T_ex-term_ cells relative to CD137^−^ CD8^+^ T cells with only a minor proportion of T_pex_ cells.

Analysis of CD8^+^ T cell clusters (Supplemental Figure 3A) from spleen and tumor demonstrated that CD137 expressing clusters 2, 7 and 10 (Supplemental Figure 3, B and C) co-expressed a large number of markers related to activation/exhaustion (TIGIT, TIM3, PD-1, CD39, CD160). CD8^+^ T cells in cluster 7, the majority of which was localized in the tumor (Supplemental Figure 3B), additionally expressed LAG-3 (Supplemental Figure 3C). Moreover, co-expression analysis confirmed the presence of a T_pex_-like cluster (TCF1^+^PD1^+^CD69^−^), which was preferentially found in the spleen (cluster 3). Also, a T_RM_-like cluster (CD69^+^CD103^+^) was detected, which was preferentially found in tumor (cluster 5) (Supplemental Figure 3C).

Altogether, de novo arising highly proliferative CD137^+^ CD8^+^ T cells exhibited an activated and exhausted-like, cytotoxic effector and least stem-like phenotype relative to the other human CD8^+^ T cell populations we identified in tumor-bearing HIS mice.

### Maintained expansion capacity and enrichment of *in vitro* tumor reactivity within an activated and exhausted-like human CD8^+^ T cell subset

To investigate tumor reactivity of CD137^+^ CD8^+^ T cells relative to the other T cell subsets, splenic T cells were expanded *ex vivo* (**Figure 3A**). Using a rapid expansion protocol (20), we observed an average expansion of >1000-fold after 14 days for all subsets. Interestingly, although terminally exhausted CD8^+^ T cells have generally been thought to contain reduced proliferative potential (51), there were no significant differences in the expansion capacity of CD137^+^ CD8^+^ T cells displaying the most exhausted-like, T_ex-term_ enriched phenotype (**Figure 2O** and Supplemental Figure 2, J and K) compared to the other CD8^+^ T cell populations from tumor-bearing HIS mice or bulk CD8^+^ T cells from naïve HIS mice (**Figure 3B**). All of the *ex vivo* expanded T cell subsets from tumor-bearing HIS mice exhibited differentiation biased to an effector memory-like phenotype (**Figure 3C**). While prior to *ex-vivo* expansion both PD-1 and CD39 were strongly increased in CD137^+^ CD8^+^ T cells (**Figure 2, O** and **N**), both markers were no longer significantly upregulated after expansion. In contrast, HLA-DR remained upregulated on CD137^+^ CD8^+^ T cells relative to CD137^−^PD1^−^ CD8^+^ and bulk CD8^+^ T cells from naïve mice (**Figure 3D**).

**Figure 3:**
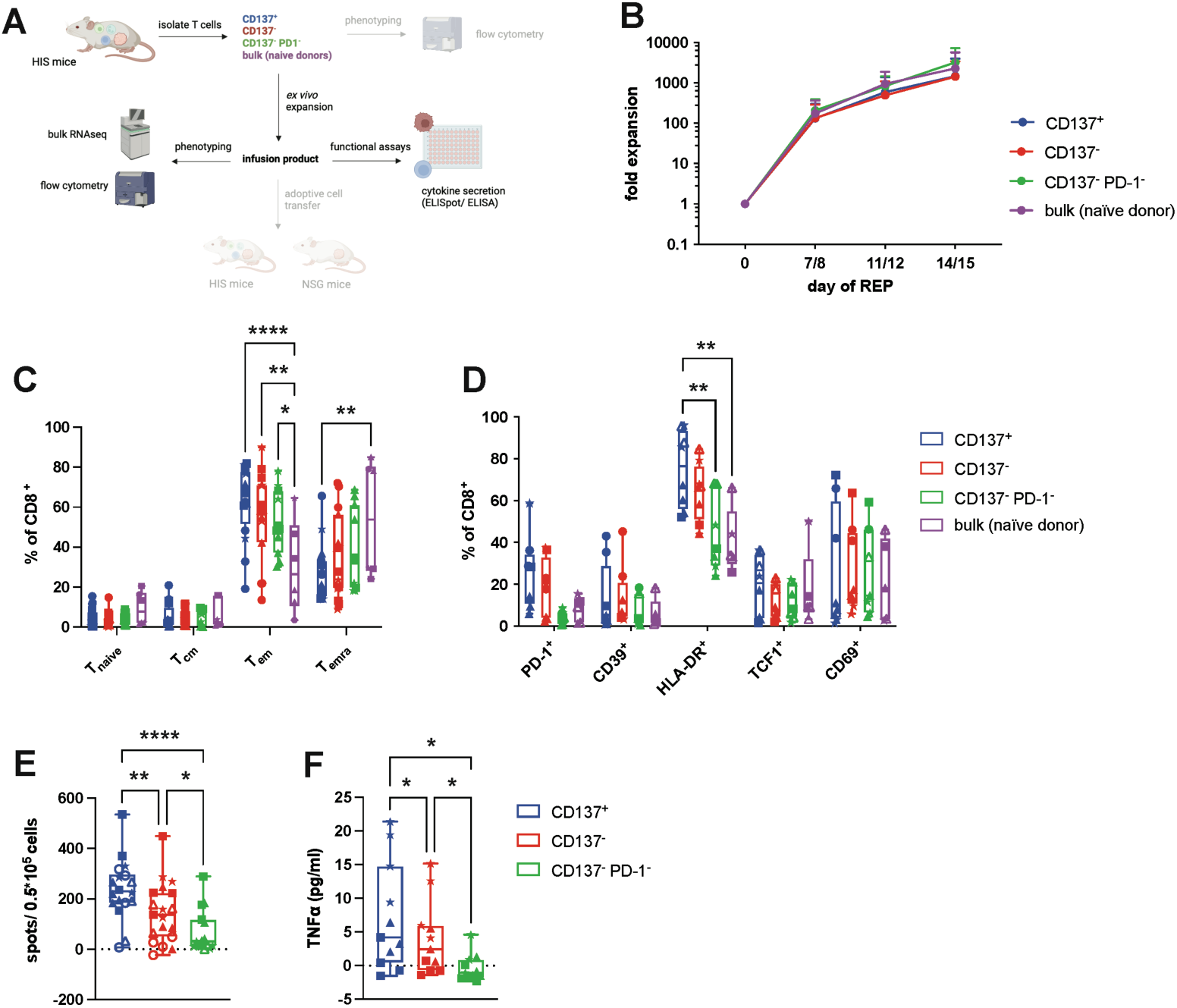
Exhausted-like human CD8^+^ T cells derived from tumor-bearing HIS mice can be expanded *ex vivo* and exhibit superior tumor-specific cytokine production. **A**, schematic of generation, expansion and characterization of T cells from HIS mice bearing autologous LCL tumors. **B**, fold expansion of FACS sorted splenic CD8^+^ T cells from tumor-bearing HIS mice (CD137^+^, CD137^−^ and CD137^−^PD1^−^) or naïve HIS mice (bulk). **C**, CD8^+^ T cell differentiation after *ex vivo* expansion defined as T_naïve_ (CD45RA^+^CD62L^+^), T_CM_ (CD45RA^−^ CD62L^+^), T_EM_ (CD45RA^−^CD62L^−^), T_EMRA_ (CD45RA^+^CD62L^−^). **D**, expression of indicated markers after *ex vivo* expansion. E, IFN-γ ELISpot of expanded T cells in 5:1 (E:T) co-culture with autologous LCL tumor cells for 24 hours. Spot count is normalized to the spots produced by expanded CD8^+^ bulk T cells from naïve HIS mice. **F**, TNF⍺ ELISA of supernatant of expanded T cells in co-culture (5:1, E:T) with autologous LCL tumor cells for 24 hours. TNF⍺ concentration is normalized to the TNF⍺ secretion from expanded CD8^+^ bulk T cells derived from naïve mice. Data are pooled from at least 3 independent experiments, n=6-19 mice per group. For each experiment, a different HPC donor was used for HIS mouse reconstitution and generation of autologous tumor. Significance by mixed-effects analysis (paired), RM one-way ANOVA or 2way ANOVA, as appropriate. Data from individual experiments are indicated by different symbols, with individual mice from the same experiment indicated by the same symbol.

After co-culture with autologous tumor cells, expanded CD137^+^ CD8^+^ T cells showed the highest IFN-γ production by ELISpot, which gradually decreased from CD137^−^ CD8^+^ T cells to CD137^−^PD-1^−^ CD8^+^ T cells (**Figure 3E**). Similarly, TNFα secretion was highest in supernatants containing CD137^+^ CD8^+^ T cells after co-culture with autologous tumor cells and lowest in CD137^−^PD-1^−^ CD8^+^ T cells (**Figure 3F**).

Together, these data indicated enrichment of tumor recognition within CD137^+^ CD8^+^ T cells that maintained *ex vivo* proliferative potential, while double negative CD137^−^PD-1^−^ CD8^+^ T cells behaved as bystander cells.

### Transcriptome analysis reveals tumor-reactive signature in CD137^+^ CD8^+^ T cells

To gain insight into differentially expressed genes (DEG) in tumor-reactive CD137^+^ CD8^+^ T cells versus CD137^−^ CD8^+^ T cells and bystander CD137^−^PD-1^−^ CD8^+^ T cells as well as bulk CD8^+^ T cells from naïve HIS mice, whole transcriptome RNA sequencing of *ex vivo* expanded T cell subsets was performed. We found a higher number of DEG for the tumor-reactive (CD137^+^) gene set versus bystander CD8^+^ T cells from tumor-bearing HIS mice (200 DEG) and versus bulk CD8^+^ T cells from naïve HIS mice (300 DEG) in comparison to the number of DEG we found for the comparison against CD137^−^ CD8^+^ T cells (53 DEG) (**Figure 4A**), the subset with the second highest tumor reactivity *in vitro* (**Figure 3, D** and **E**). Principal component analysis (PCA) was consistent with functional data of tumor-specific cytokine production (**Figure 3, D** and **E**), showing that the 4 subsets clustered in a gradual way from tumor-reactive (CD137^+^) to less tumor-reactive (CD137^−^) and bystander (CD137^−^PD-1^−^) CD8^+^ T cells and finally to bulk CD8^+^ T cells from naïve HIS mice, with the biggest overlap between CD137^−^PD-1^−^ bystander CD8^+^ T cells and bulk human CD8^+^ T cells from naïve HIS mice (**Figure 4B**).

**Figure 4:**
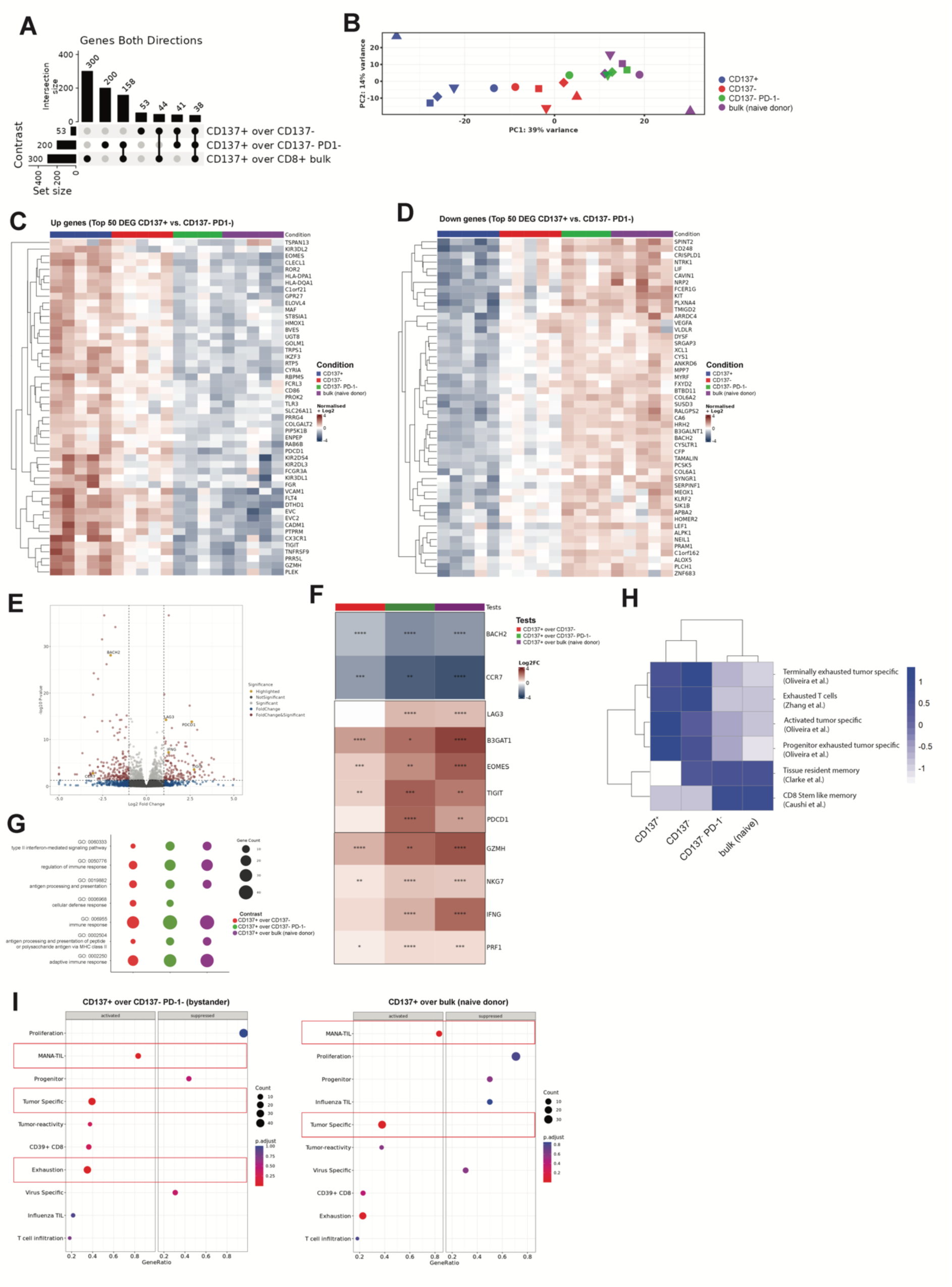
Transcriptomic profiling of tumor-reactive CD137^+^ CD8^+^ T cells. Transcriptome analysis and pathway analysis of expanded CD8^+^ T cells from tumor-bearing HIS mice (CD137^+^, CD137^−^ and CD137^−^PD1^−^) or naïve HIS mice (bulk). **A,** Upset plot (intersect) showing **number** of differentially expressed genes between groups in bulk RNAseq. p(FDR) < 0.05, log2 FC > 1.5. **B**, PCA plot of RNAseq showing PC1 and PC2. **C,** Top 50 upregulated and **D,** downregulated genes of T cell subsets based on the DEG between CD137^+^ vs. CD137^−^PD1^−^ CD8^+^ T cells. p(FDR) < 0.05, log2 FC > 1.5. **E,** Volcano plot showing DEG between CD137^+^ and CD137^−^PD1^−^ CD8^+^ T cells with genes of interest highlighted in yellow. **F**, differential expression of genes of interest between groups. **G**, Overrepresentation analysis (ORA) of upregulated pathways in CD137^+^ CD8^+^ T cells. **H**, Gene module score analysis of published signatures described on the right. **I**, Gene set enrichment analysis (GSEA) of signatures described on the y-axis. Gene ratio (# genes related to GO term / total number of sig genes) is displayed on the x-axis. Signatures with an adjusted p-value <0.05 are highlighted with a red box. Shown are data from 4-5 individual experiments, each experiment with different human HPC donor for reconstitution of HIS mice and autologous tumor.

The transcriptomic profile that we found in tumor-reactive CD137^+^ CD8^+^ T cells suggested an exhausted/activated-like phenotype with increased expression of *TNFRSF9*, *LAG3*, *PDCD1, TIGIT* and *HLA-DPA1/HLA-DQA1* (**Figure 4, C, E and F**). We also observed enhanced expression of effector molecules associated with cytotoxicity (*GZMH, NKG7* and *PRF1*) as well as *IFNG* (**Figure 4, C, E and F**). Expression of the naïve or stem-like genes *CCR7*, *LEF1* and *BACH2* were diminished (**Figure 4, D-F**) while the senescence marker CD57 (52) (*B3GAT1*) and the transcription factor *EOMES* showed the highest transcript level in the CD137^+^ subset (**Figure 4, C and F**). Although EOMES has been shown to be involved in T cell exhaustion, it has also been reported to be commonly co-expressed with effector molecules in effector T cells and to be essential for anticancer T cell function (53,54).

Among upregulated Gene Ontology (GO) terms comparing tumor-reactive CD137^+^ CD8^+^ T cells against the other three populations, we found the terms interferon-gamma mediated signaling (GO:0060333), antigen processing and presentation (GO:0019882), antigen processing and presentation of peptide or polysaccharide antigen via MHC class II (GO:0002504), immune response (GO:0006955) and adaptive immune response (GO:0002250) (**Figure 4G**). Upregulation of these pathways in the CD137^+^ population indicated stronger T cell activation and response in comparison to the other populations.

To determine T cell states after *ex vivo* expansion, we compared our transcriptome data to published datasets of different T cell states from human cancers (31,46,55–59). A high overlap of signatures originally found in tumor-specific CD8^+^ T cells from human cancers describing terminally exhausted, exhausted, activated, and progenitor exhausted T cell states was found for CD137^+^ CD8^+^ T cells (**Figure 4H**). Overlap with these different signatures might hint to a degree of heterogeneity within the CD137^+^ CD8^+^ T cell population after *ex vivo* expansion but confirmed the overall activated/exhausted-like state of these cells that we already observed before *ex vivo* expansion. Signatures for tissue-resident memory and CD8^+^ stem-like memory were downregulated in the CD137^+^ CD8^+^ population. Strikingly, comparing tumor-reactive CD137^+^ CD8^+^ T cells to bystander CD137^−^PD-1^−^ CD8^+^ T cells or bulk CD8^+^ T cells from naïve HIS mice, gene sets that have been described to define tumor-reactive T cells in patients with cancer were significantly upregulated (‘MANA TIL’ (58), ‘tumor specific’ (56)) (**Figure 4I**). These results suggested that the antitumor T cell response in autologous tumor-bearing HIS mice mirrors relevant aspects of the T cell response in cancer patients, thus further supporting the potential translational value of our model.

### Correlation of clonal expansion with effector function

As another surrogate for tumor recognition, we analyzed clonality of the CD8^+^ T cell receptor (TCR) repertoire, anticipating tumor-reactive CD8^+^ T cells would have higher clonal expansion and thus lower diversity than bystander CD8^+^ T cells. Indeed, we found the total number of clonotypes tending to be the smallest in the tumor-reactive CD137^+^ subset, and to gradually increase from the less tumor-reactive CD137^−^ subset to CD137^−^PD-1^−^ bystander CD8^+^ T cells to bulk CD8^+^ T cells from naïve HIS mice (**Figure 5A**), following the gradient of tumor recognition we observed by tumor-specific cytokine production *in vitro* (**Figure 3, D** and **E**). Likewise, the estimated sample diversity for the tumor-reactive CD137^+^ subset was lower than for the other populations (**Figure 5B** and Supplemental Figure 4A). In line with this, the CD137^+^ population had a reduced abundance of rare or very small clonotypes (**Figure 5C** and Supplemental Figure 4, B and C). Conversely, tumor-reactive CD137^+^ CD8^+^ T cells from tumor-bearing HIS mice demonstrated a significantly higher abundance of hyperexpanded clones (**Figure 5D**) with the 20 biggest clones occupying a significantly larger repertoire space than in the other populations, suggesting polyclonal expansion (Supplemental Figure 4B).

**Figure 5:**
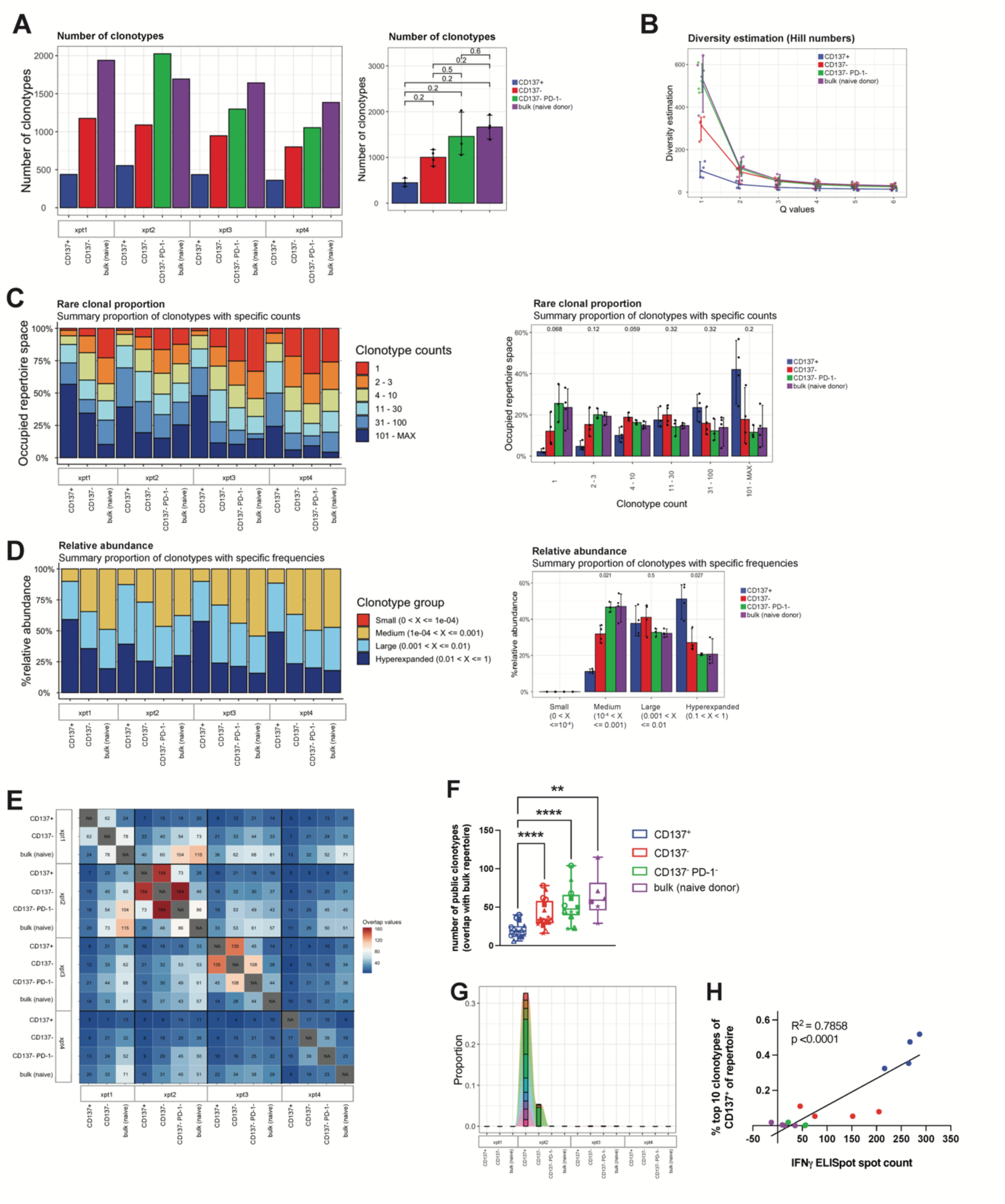
TCR profiling of tumor-reactive CD137^+^ CD8^+^ T cells. **A,** total number of individual clonotypes found per population. Left: data from individual experiments, right: pooled data for group analysis. Wilcoxon test. **B,** TCR (CDR3 of *TRA*, *TRB*, *TRG* and *TRD*) sequence sample diversity estimation using Hill numbers method, with Q=1 describing the Shannon diversity. **C,** rare clonal proportion showing the occupied repertoire space by clonotypes with defined counts (1, 2-3, 4-10, etc.). Left: data from individual experiments, right: pooled data for group analysis. Wilcoxon test. **D,** relative abundance of clonotypes with defined frequencies (size). Left: data from individual experiments, right: pooled data for group analysis. Wilcoxon test. **E,** repertoire overlap analysis cross-comparing every population from every experiment (each with different donor). **F**, repertoire overlap comparing the repertoire of bulk CD8^+^ T cells from naïve HIS mice to the populations from tumor-bearing HIS mice from individual experiments (each with different donor; data from individual experiments are indicated by different symbols). Mixed effects analysis with Tukey’s multiple comparisons test. **G**, tracking of clonotypes over populations. The top 10 most abundant clonotypes of the TCR repertoire of CD137^+^ CD8^+^ T cells from one representative experiment are shown. **H**, proportion of the top 10 most abundant clonotypes (from repertoires of CD137^+^ CD8^+^ T cells) in the repertoire of all populations, correlated with the spot count of IFN-γ ELISpot. Shown are data from 4 individual experiments, each experiment with different human HPC donor for reconstitution of HIS mice and autologous tumor.

To track the overlap of clonotypes between individual experiments (i.e. donor dependency) and populations, we next performed TCR repertoire overlap analysis (**Figure 5E**). We found that the tumor-reactive CD137^+^ population showed the biggest overlap with the CD137^−^ subset and only to a minor extend an overlap with CD137^−^PD-1^−^ bystander CD8^+^ T cells or bulk CD8^+^ T cells from naïve HIS mice (**Figure 5, E and G**). Also, analyzing the repertoire of the bulk naïve CD8^+^ T cell population for each experiment by cross-comparing to each population from each experiment, the overlap with the CD137^+^ population was significantly less than the overlap with every other population (**Fig. 5F**). These findings suggested that clonotypes arising due to the conditions of expansion were less present in the CD137^+^ population and that other, presumably tumor-reactive, clonotypes expanded.

In addition, we observed that the expansion of individual bona fide tumor-reactive clonotypes was highly donor specific. When comparing the 10 most abundant clonotypes of the CD137^+^ population per experiment (i.e. per donor), almost no overlap with the CD137^+^ TCR repertoire of any other experiment was detected (**Figure 5G** and Supplemental Figure 4D). Finally, to explore an association of clonal expansion with functionality, we correlated the abundance of the top 10 bona fide tumor-reactive clonotypes of the CD137^+^ population with *in vitro* tumor reactivity and found a highly significant correlation of clone size abundance and tumor-specific IFN-γ production by CD8^+^ T cells (**Figure 5H**). Overall, these data indicated that CD137^+^ CD8^+^ T cells derived from tumor-bearing HIS mice expanded polyclonally in a human donor- and tumor-specific manner.

### Human CD8^+^ T cells with an activated and exhausted-like phenotype reduce tumor growth *in vivo*

We next performed adoptive cell transfer (ACT) experiments of *ex vivo* expanded CD8^+^ T cells in NSG mice previously engrafted with autologous tumors (**Figure 6A**). We initially selected NSG mice as recipients to unambiguously track the transferred T cells and to be able to distinguish their anticancer activity from the endogenous antitumor T cell response in HIS mice. In NSG mice, only ACT with expanded CD137^+^ CD8^+^ T cells resulted in reduced tumor growth in a substantial number of recipients, while transfer of bystander CD8^+^ T cells from tumor-bearing HIS mice or bulk CD8^+^ T cells from naïve HIS mice did not (**Figure 6, B-D**). Specifically, adoptive transfer of human CD137^+^ CD8^+^ T cells from tumor-bearing HIS donors resulted in a partial response (≥30% tumor reduction compared to ‘tumor only’ group) in almost half of recipients (6 out of 13 tumor-bearing NSG recipients, 46%), while this was observed in only 13% of NSG mice treated with CD137^−^PD-1^−^ CD8^+^ T cells from tumor-bearing HIS mice (1/8) and no such response (0%) was seen in NSG recipients of bulk CD8^+^ T cells from naïve HIS mice (0/6) (**Figure 6D**).

**Figure 6:**
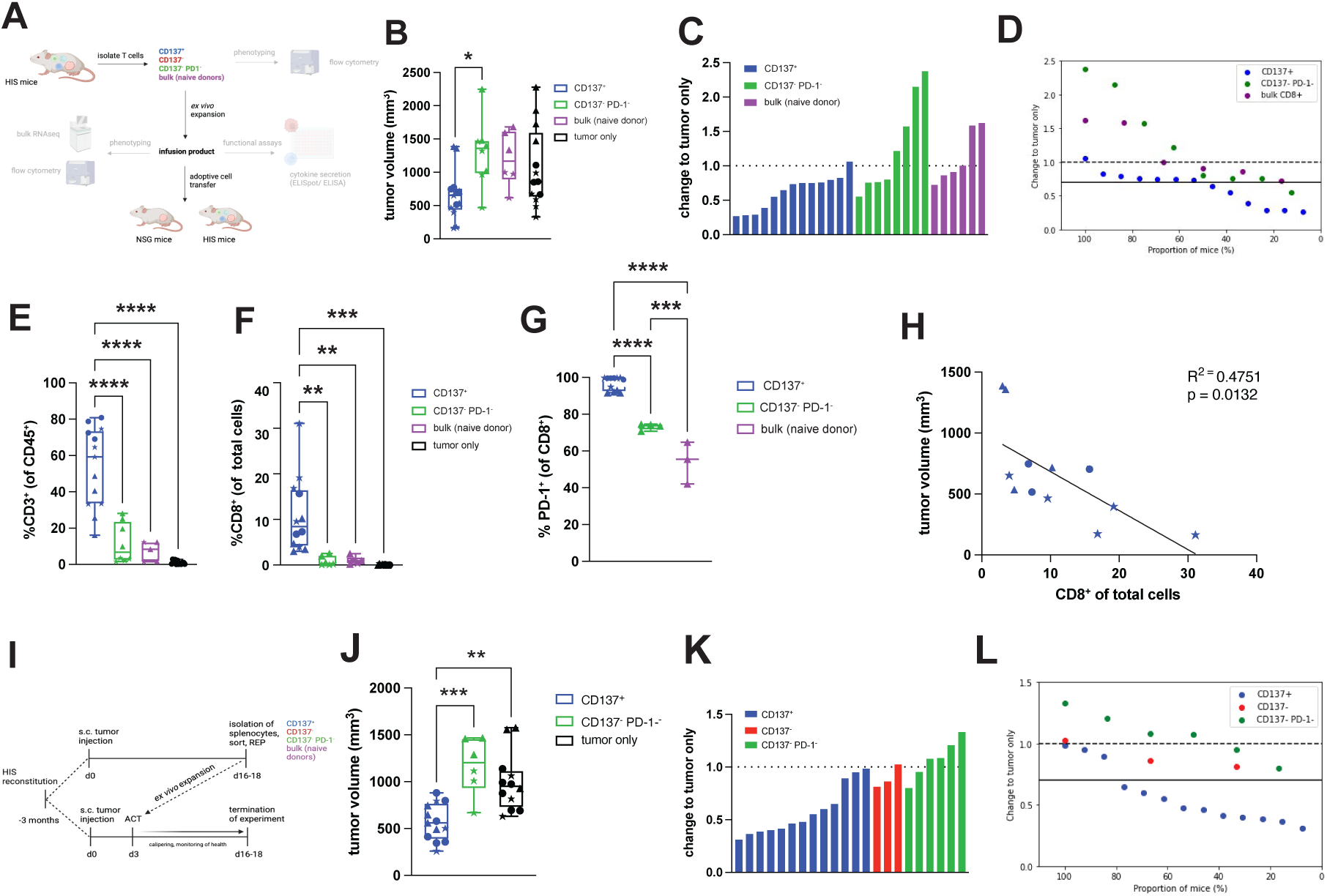
CD137^+^ CD8^+^ T cells from tumor-bearing HIS mice show superior anticancer activity in NSG and HIS recipients bearing autologous tumors. **A,** schematic of generation of tumor-reactive T cells and subsequent ACT. NSG mice were injected with 2 × 10^6^ LCL s.c. in the flank and after three days, 10 × 10^6^ *ex vivo* expanded T cells were adoptively transferred intravenously. Transferred T cells and LCL tumors were autologous to each other. **B**, tumor volume on the day of sacrifice in NSG recipient mice after ACT of the indicated cell populations. **C,** waterfall plot of tumor size in NSG recipient mice of ACT on the day of sacrifice relative to the tumor volume of control mice (no ACT). Bars depict individual mice. **D**, proportion of NSG recipient mice showing partial response (≥ 30% tumor reduction compared to control NSG mice) after ACT of the indicated cell population. **E**, frequency of CD3^+^ T cells (% of human CD45^+^ cells) in TIL of NSG mice after ACT of the indicated cell populations, measured by flow cytometry. **F**, frequency of CD8^+^ T cells (% of total cells) in tumors of NSG mice after ACT of the indicated cell populations, measured by immunohistochemistry. **G**, frequency of PD-1 expression on CD8^+^ T cells in TIL of NSG mice after ACT of the indicated cell populations. **H,** correlation between tumor volume and infiltration of CD8^+^ T cells (measured by IHC) in tumors of NSG mice after adoptive transfer of CD137^+^ CD8^+^ T cells. **I**, schematic of ACT. CD137^+^ CD8^+^ T cells, CD137^−^ CD8^+^ T cells and CD137^−^ PD-1^−^ CD8^+^ T cells were isolated from spleen of tumor-bearing HIS mice or bulk CD8^+^ T cells from spleen of naïve HIS mice and expanded *ex vivo*. Recipient HIS mice were injected with 2 × 10^6^ LCL s.c. in the flank and after three days, *ex vivo* expanded T cells were adoptively transferred intravenously. Tumor-bearing HIS recipient mice received 2 × 10^6^ T cells without prior conditioning/lymphodepletion. Donor and recipient HIS mice as well as LCL were autologous to each other. **J,** tumor volume on the day of sacrifice in HIS recipient mice after ACT of the indicated cell populations. **K,** waterfall plot of tumor size on the day of sacrifice of HIS mice receiving ACT relative to the tumor volume of control HIS mice (no ACT). Bars depict individual mice. **L**, proportion of HIS recipient mice showing partial response (≥ 30% tumor reduction compared to control HIS mice) after ACT of the indicated cell population. Data are pooled from 1-3 independent experiments, n=3-13 per group. For each experiment, a different HPC donor was used for HIS mouse reconstitution and generation of autologous tumor. Significance by one-way ANOVA. Data from individual experiments are indicated by different symbols, with individual mice from the same experiment indicated by the same symbol.

Furthermore, a higher frequency of T cells was found in tumor (**Figure 6, E and F**), blood (Supplemental Figure 5A) and spleen (Supplemental Figure 5B) of tumor-bearing NSG mice receiving CD137^+^ CD8^+^ T cells compared to recipients transferred with any of the other cell populations. Flow cytometric analysis of tumor-infiltrating lymphocytes (TIL) revealed high CD3^+^ T cell frequencies (53.65%±21.94) after ACT of expanded CD137^+^ CD8^+^ T cells, but only 11.64%±11.0 after transfer of bystander CD8^+^ T cells and 7.25%±5.38 after transfer of bulk CD8^+^ T cells from naïve HIS mice (**Figure 6E**). Likewise, immunohistochemistry of tumors of NSG mice confirmed T cell infiltration selectively after ACT with CD137^+^ CD8^+^ T cells (**Figure 6F**).

Interestingly, while infusion products mainly contained a mix of T_EM_-like and T_EMRA_ CD8^+^ T cells (Figure 3C), almost exclusively T_EM_-like cells were recovered from blood (Supplemental Figure 5C), spleen (Supplemental Figure 5D) and TIL (Supplemental Figure 5E) of recipient NSG mice. Of note, whereas PD-1 was highly expressed on splenic CD137^+^ CD8^+^ T cells in tumor-bearing HIS mice (96.49%±3.1) (**Figure 2N**), PD-1 expression on this population decreased during antigen-independent *ex vivo* expansion (32.45%±29.17) (Supplemental Figure 5F). However, after adoptive transfer of expanded CD137^+^ CD8^+^ T cells into tumor-bearing NSG mice, PD-1 expression among CD8^+^ T cells increased again and was significantly higher in TIL (96.7%±3.65) (**Figure 6G**), blood (Supplemental Figure 5G) and spleen (Supplemental Figure 5H) compared to NSG mice transferred with bystander CD8^+^ T cells or bulk CD8^+^ T cells. Apart from the superior tumor control, these data further indicated enrichment of tumor reactivity within CD137^+^ CD8^+^ T cells, with very high PD-1 expression serving as a potential marker for productive tumor recognition, but not loss of effector function. As another index of the anticancer activity of CD137^+^ CD8^+^ T cells, the degree of tumor infiltration with CD8^+^ T cells after ACT was inversely correlated with tumor size (**Figure 6H**). Thus, the higher frequency of CD8^+^ T cells in tumors was likely driven by increased tumor recognition of transferred CD137^+^ CD8^+^ T cells, and these cells mediated tumor cell killing.

Lastly, ACT of tumor-reactive T cells into autologous tumor-bearing HIS recipient mice (**Figure 6I**) presented an opportunity to model polyclonal TIL transfer therapy in cancer patients (60) in an immunocompetent humanized tumor setting. As observed with NSG recipients, transfer of polyclonal tumor-reactive CD137^+^ CD8^+^ T cells into autologous tumor-bearing HIS recipient mice reduced tumor growth (**Figure 6, J-L**). A partial response (≥30% tumor reduction compared to ‘tumor only’ group) was observed in 77% of HIS mice (10/13) receiving adoptive transfer of autologous human CD137^+^ CD8^+^ T cell derived from tumor-bearing HIS donors, while no such response (0%) was observed in any of the HIS mice (0/9) adoptively transferred with autologous CD137^−^ or CD137^−^PD-1^−^ CD8^+^ T cells derived from tumor-bearing HIS donors (**Figure 6L**). Importantly, this was achieved without prior conditioning of HIS recipient mice or cytokine support, and the number of transferred tumor-reactive CD137^+^ CD8^+^ T cells was fivefold lower compared with transfer in NSG recipients. These data established superior anticancer activity of CD137^+^ CD8^+^ T cells with an activated and exhausted-like phenotype in the most complete humanized setting.

Altogether, these adoptive transfer experiments confirmed anticancer activity of human CD137^+^ CD8^+^ T cells that developed in autologous tumor-bearing HIS mice, suggesting that our model captured relevant aspects of the immunobiology of human antitumor T cell responses.

## Discussion

Using in-depth phenotypic analyses, we established the presence of major, widely recognized human CD8^+^ T cell subsets known to be important in tumor immunology, like progenitor exhausted (T_pex_; TCF1^+^PD-1^+^), terminally exhausted (T_ex-term_; TCF1^−^PD-1^+^CD69^+^) or tissue resident-like (T_RM_; CD69^+^CD103^+^) T cells in our autologous humanized tumor model. Interestingly, we found CD8^+^ T_pex_ cells to be more abundant in secondary lymphoid tissue (spleen) compared to tumor and vice versa CD8^+^ T_ex-term_ cells to be more abundant in tumor compared to spleen. This seems consistent with the mechanisms of formation of antitumor immunity as currently understood (61). Likewise, tissue-resident-like (T_RM_) T cells were enriched in the tumor relative to spleen, as would be expected. Furthermore, we identified a proliferative human CD8^+^ T cell subset in tumor-bearing HIS mice that expressed activation and exhaustion markers including very high levels of PD-1 and CD39 while having low expression of the progenitor marker TCF1 but high expression of TOX. This CD137^+^ CD8^+^ T cell subset, while being PD-1^hi^CD39^+^, displayed increased abundance of a T_ex-term_ phenotype with co-expression of inhibitory receptors like TIM-3, TIGIT, CD160 and galectin-9. At the same time, CD137^+^ CD8^+^ T cell contained fewer T_RM_-like cells, which was reflected by a relative lack of a tissue-resident signature even after expansion. However, we note that the expansion, which was necessary to achieve the cell numbers required for adoptive transfer, may have led to some alteration of populations, as evidenced, for example, in the downregulation of PD-1 and CD39 on the protein level after expansion. Nevertheless, most activation and exhaustion markers were still increased on the transcript level in expanded CD137^+^ CD8^+^ T cells relative to the other populations. Furthermore, expanded CD137^+^ CD8^+^ T cells from tumor-bearing HIS mice showed enrichment in gene signatures of exhaustion. Functionally, CD137^+^ CD8^+^ T cells exhibited increased tumor reactivity as demonstrated by enhanced cytokine production after coculture with autologous tumor cells. Importantly, by means of adoptive transfer, we uniquely provide direct *in vivo* evidence of superior anticancer activity of these human tumor-reactive effector CD8^+^ T cells displaying an activated and exhausted-like phenotypic state.

We have leveraged this human CD8^+^ T cell subset arising de novo in tumor-bearing HIS mice for transcriptomic analysis and TCR profiling to demonstrate across donors overlap with tumor-reactive CD8^+^ T cell signatures from human cancers (56,58) as well as tumor-induced polyclonal expansion. Polyclonality of neoantigen-specific CD8^+^ T cells in blood and tumor has recently been shown to be associated with response to PD-1 immunotherapy in patients with melanoma (16). Congruence of our model data across donors with these clinical benchmarks suggests robustness and translational validity of our findings.

Of note, CD39^+^CD8^+^ T cells expressing high levels of PD-1 have been described to be enriched for tumor reactivity and clonal proliferation, predicting efficacy of immunotherapy in human lung cancer (46). Likewise, the abundance of intratumoral CD39^+^CD8^+^ T cells displaying upregulated exhaustion markers has been shown to be prognostic for improved progression-free and overall survival in individuals with treatment-naïve early stage lung cancer (62). Furthermore, PD-1^hi^CD39^+^ CD8^+^ T cells correlated with improved survival in patients with endometrial cancer (45) and the signature of similar CD8^+^ T cells with increased TOX expression predicted survival in patients with breast cancer (44). Recently, pre-existing exhausted-like tumor-reactive CD8^+^ T cells have been associated with favorable outcome to immunotherapy in head and neck cancer (63). On the other hand, higher abundance of TCF1^+^ stem-like CD8^+^ T cells with a CD39^−^CD69^−^ phenotype in TIL products was associated with better response after TIL transfer in patients with melanoma (64). However, by single-cell analysis of lung tumors, this phenotype could not be identified in neoantigen-reactive T cells (65). While we have not addressed and thus cannot exclude a contribution of the minor TCF1^+^ fraction within the human tumor-reactive CD8^+^ T cells in our model, our data emphasize the key importance of exhausted-like effector CD8^+^ T cells in the antitumor immune response.

The striking phenotypic similarity of tumor-reactive CD8^+^ T cells that have been shown to correlate with clinical benefit and the polyclonally expanded human tumor-reactive CD8^+^ T cell cells from tumor-bearing HIS mice that we demonstrated to possess superior *in vivo* anticancer activity highlights the translational potential of our model. Hence, the novel immunocompetent human tumor model that we developed using HIS mice with human immune cells reconstituted from hematopoietic progenitor cells that are autologous to tumor might bridge a gap between syngeneic mouse tumor models and human-specific antitumor immune responses (13,66). Our model demonstrates hallmarks of human antitumor T cell responses and thus may provide a tool that should allow further investigation of drivers of anticancer activity in human CD8^+^ T cells in cancer immunotherapy research. Overall, the endogenous human immune system in HIS mice markedly influenced tumor and host, allowing for experimental interrogation of the interfaces of human tumor and autologous human antitumor immunity in a small animal model. Together, fully activated effector and exhausted-like human CD8^+^ T cells directly mediate anticancer activity. Our data support clinical exploration of exhausted-like effector CD8^+^ T cell subsets alongside the recently more favored stem-cell like phenotype for use in adoptive transfer and engineered T cell therapies.

