## Supplemental Figures for "Human effector CD8^+^ T cells with an exhausted-like phenotype control tumor growth *in vivo* in a humanized tumor model"

#### Supplementary Material

**Supplemental Figures 1-5**

### Human effector CD8$^+$ T cells with an exhaustion-like phenotype control tumor growth *in vivo* in a humanized tumor model

Juliane Mietz[1], Meike Kaulfuss[1], Lukas Egli[1], Lennart Opitz[2], Christian Münz[3], Obinna Chijioke[1,4]

[1]Cellular Immunotherapy, Institute of Experimental Immunology, University of Zürich, Zürich, Switzerland
[2]Functional Genomics Center Zürich, University of Zürich/ETH Zürich, Zürich, Switzerland
[3]Viral Immunobiology, Institute of Experimental Immunology, University of Zürich, Zürich, Switzerland
[4]Institute of Medical Genetics and Pathology, University Hospital Basel, Basel, Switzerland

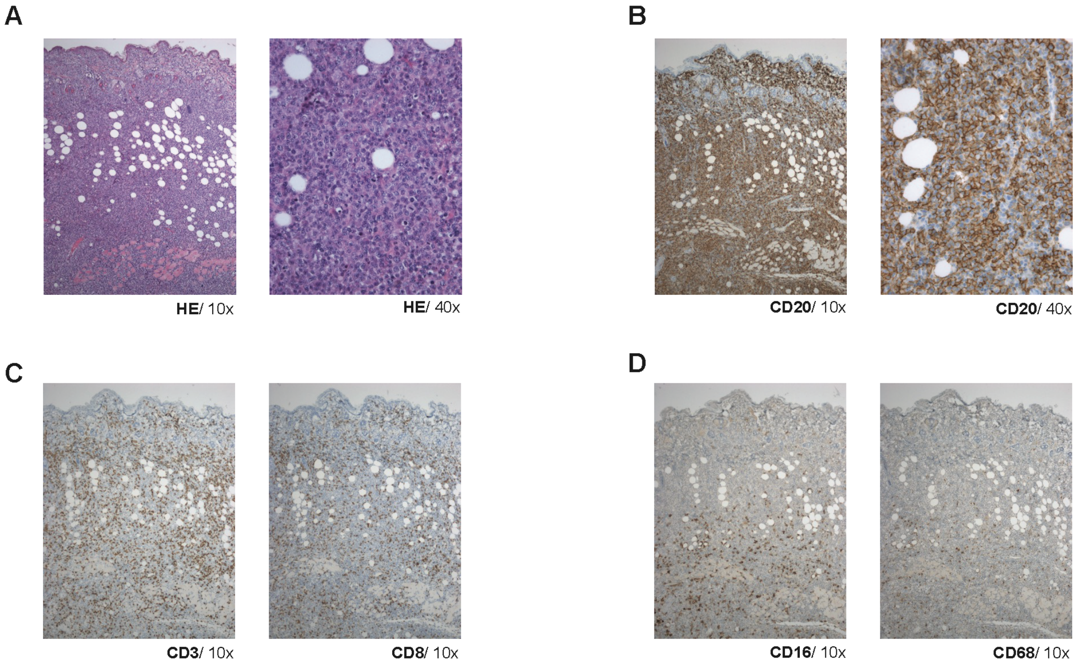


**Supplemental Figure 1: Tumor infiltration by autologous human immune cells in HIS mice.** Tumor histology by HE staining (**A**) and immunohistochemistry (IHC) showing CD20 expression of tumor cells (**B**). Human immune cell infiltration by IHC on tumor sections with staining for CD3 (left) and CD8 (right) (**C**) and CD16 (left) and CD68 (right) (**D**).

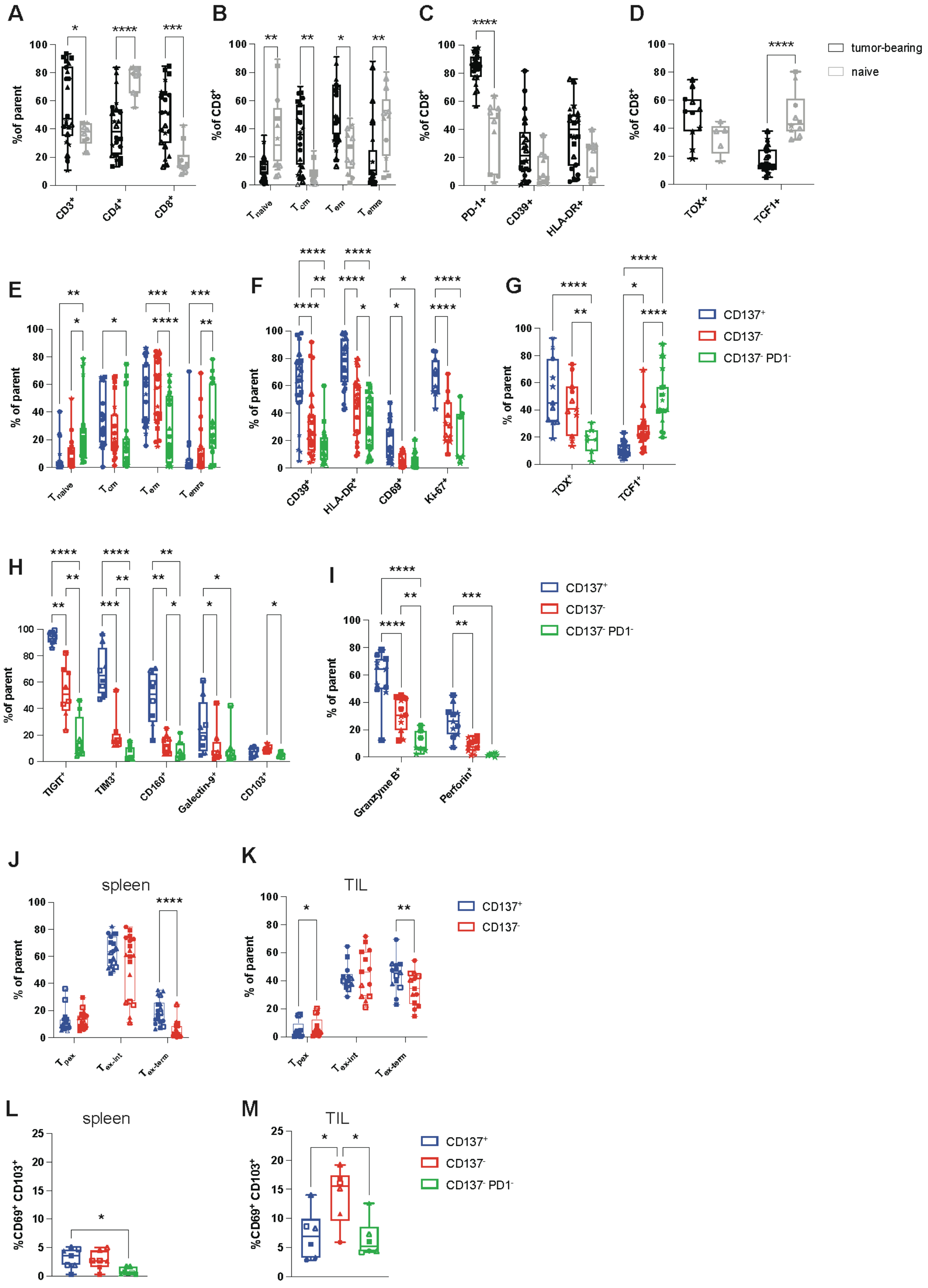


**Supplemental Figure 2: T cell profiling in peripheral blood, spleen and tumor of tumor-bearing HIS mice.** PBMC, splenocytes and TIL were isolated from tumor-bearing HIS mice and analyzed by flow cytometry. **A,** frequency of T cells in peripheral blood of tumor-bearing or non-tumor bearing HIS mice (naïve); parent population refers to frequency (%) of $CD3^+$ T cells within human $CD45^+$ cells and $CD4^+$ and $CD8^+$ T cells within $CD3^+$ T cells. **B**, $CD8^+$ T cell differentiation in peripheral blood defined as $T_{naïve}$ ($CD45RA^+CD62L^+$), $T_{CM}$ ($CD45RA^-CD62L^+$), $T_{EM}$ ($CD45RA^-CD62L^-$), $T_{EMRA}$ ($CD45RA^+CD62L^-$) in tumor-bearing or naïve HIS mice. **C-D**, expression of indicated markers on human $CD8^+$ T cells in peripheral blood of tumor-bearing or naïve HIS mice. **E-I**, proportion of $CD8^+$ T cell subsets and expression of individual markers in spleen of tumor-bearing HIS mice within $CD137^+$, $CD137^-$ or $CD137^-PD\text{-}1^-$ populations. (J-K), distribution of $T_{pex}$ ($TCF1^+PD\text{-}1^+$), $T_{ex\text{-}int}$ ($TCF1^-PD\text{-}1^+CD69^-$) and $T_{ex\text{-}term}$ ($TCF1^-PD\text{-}1^+CD69^+$) subsets within $CD137^+$ or $CD137^-$ populations from spleen (**J**) or tumor (TIL) (**K**) of tumor-bearing HIS mice. (L-M), $T_{RM}$-like cells ($CD69^+CD103^+$) within $CD137^+$, $CD137^-$ or $CD137^-PD\text{-}1^-$ $CD8^+$ T cell populations in spleen (**L**) or tumor (TIL) (**M**) of tumor-bearing HIS mice. Data are pooled from 2-6 individual experiments, n=7-23. For each experiment, a different HPC donor was used for HIS mouse reconstitution and generation of autologous tumor. Significance by one-way ANOVA or 2way ANOVA, as appropriate. Data from individual experiments are indicated by different symbols, with individual mice from the same experiment indicated by the same symbol.

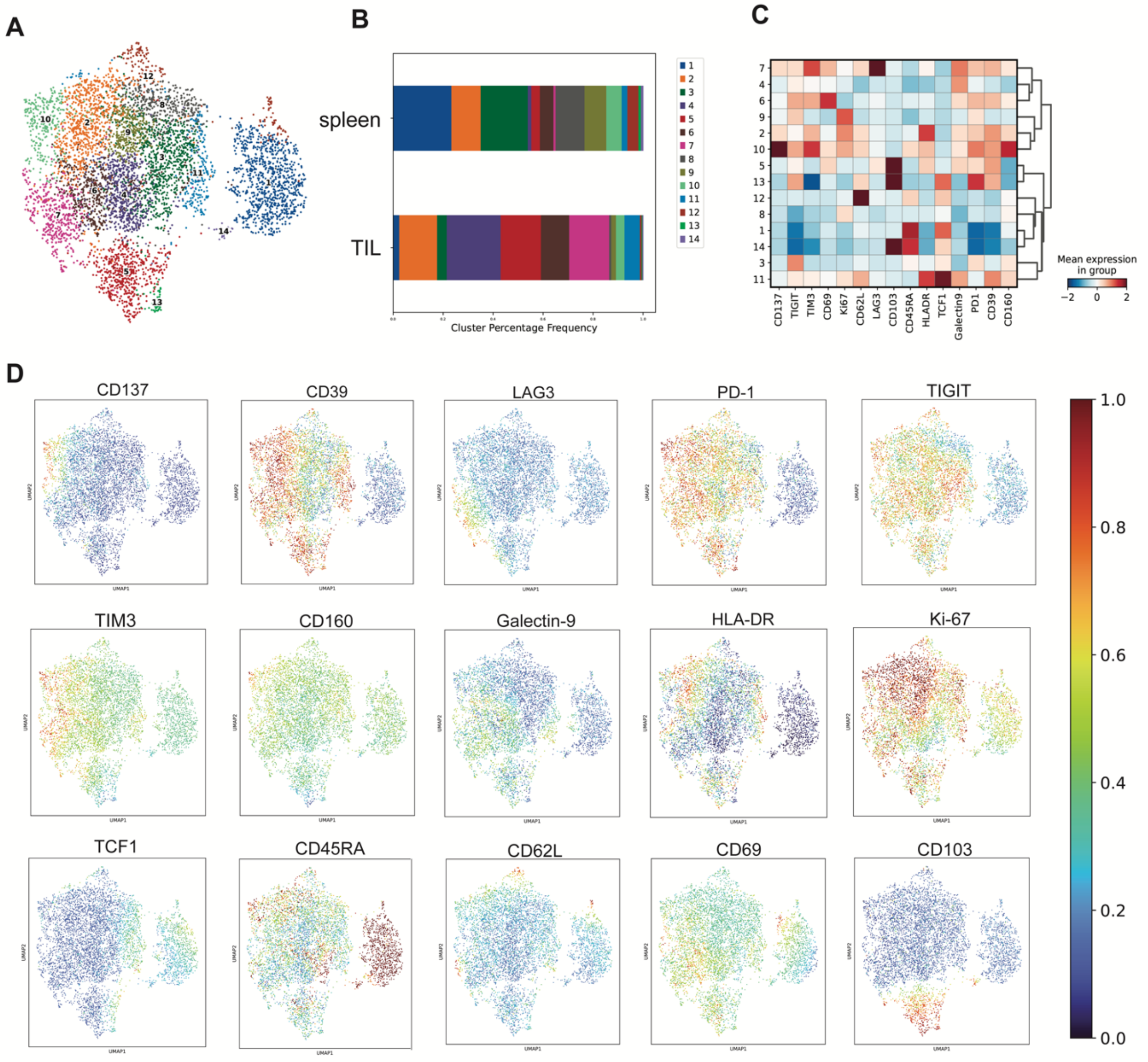


**Supplemental Figure 3: Co-expression analysis of CD8$^+$ T cells in tumor-bearing HIS mice.** Splenocytes and TIL were isolated from tumor-bearing HIS mice and CD8$^+$ T cells analyzed by flow cytometry. **A**, cluster visualization by UMAP of CD8$^+$ T cells from spleen and TIL. **B**, bar chart showing distribution of CD8$^+$ T cell clusters in spleen and TIL. **C**, heatmap depicting mean expression of individual markers in the respective CD8$^+$ T cell clusters. **D**, UMAP plots showing expression of individual markers. Data are pooled from 4 independent experiments, n=6-8 per group. For each experiment, a different HPC donor was used for HIS mouse reconstitution and generation of autologous tumor.

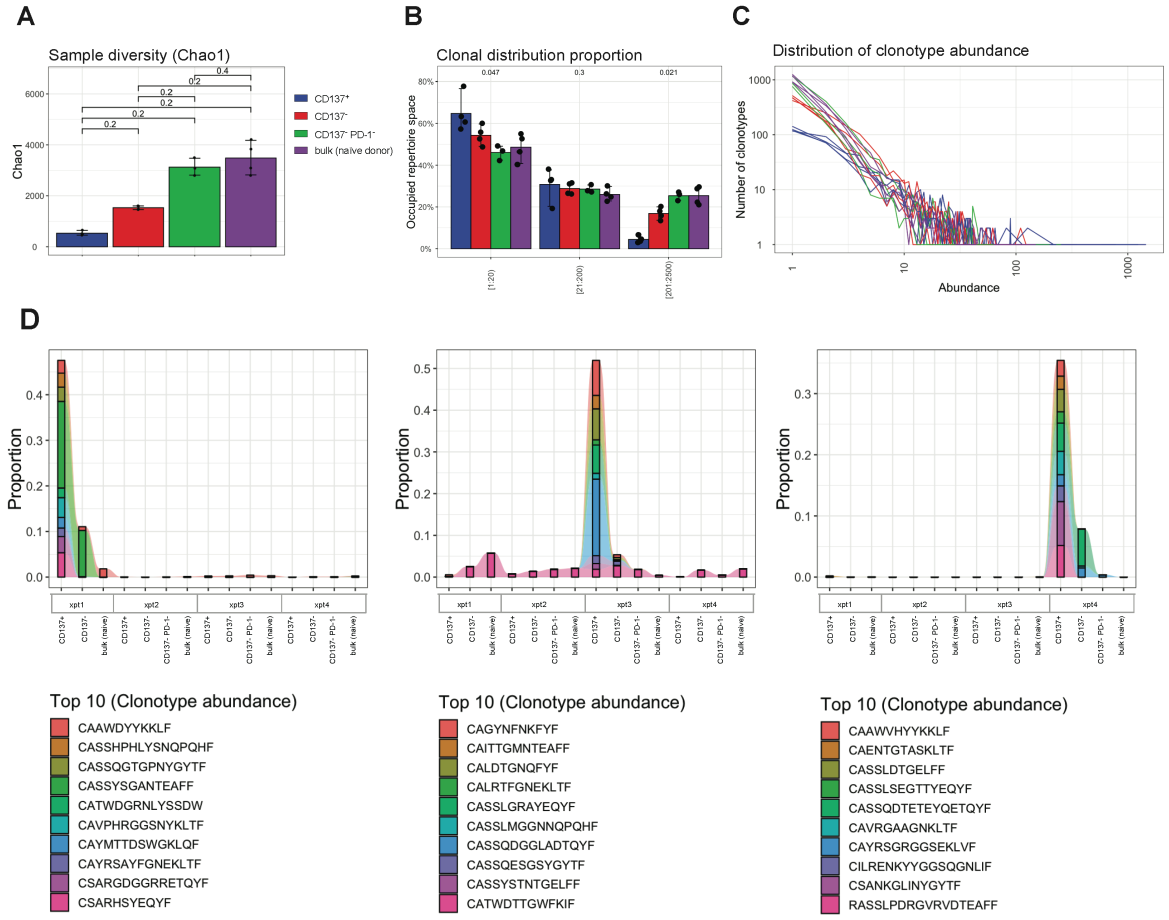


**Supplemental Figure 4: Additional TCR repertoire analyses including clonotype tracking of human tumor-reactive CD137$^+$ CD8$^+$ T cells. A,** TCR sequence (CDR3 of *TRA*, *TRB*, *TRG* and *TRD*) sample diversity estimation using Chao1 method. Wilcoxon test. **B,** clonal proportion distribution of clonotypes with the indicated indices on the x-axis (e.g. [1:20) contains the 20 largest clonotypes). **C**, distribution of clonotype abundance. **D,** tracking of clonotypes over populations and experiments/donors. The top 10 most abundant clonotypes of the TCR repertoire of the CD137$^+$ population from 3 independent experiments are shown (each experiment with different human HPC donor for reconstitution of HIS mice and autologous tumor).

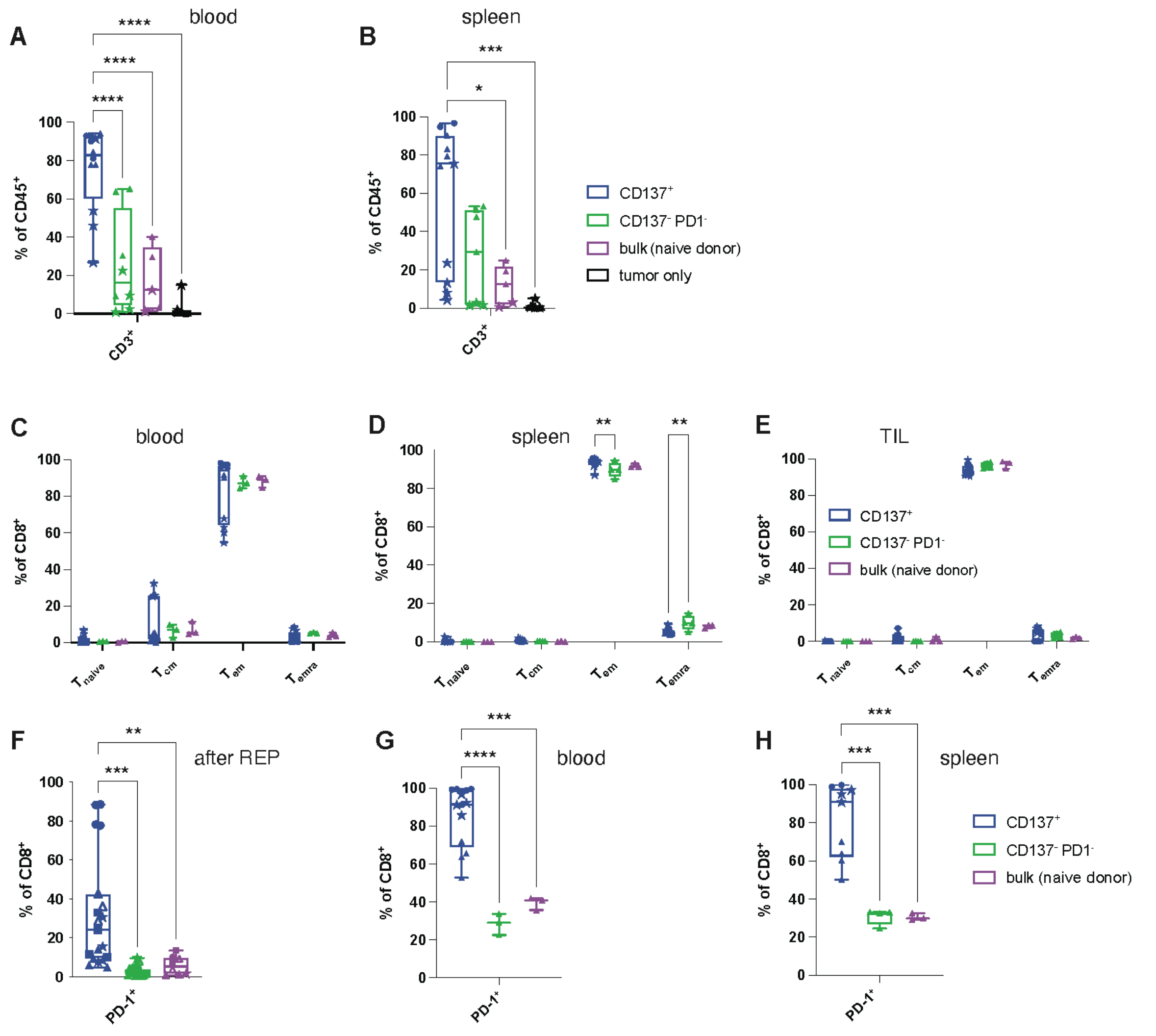


**Supplemental Figure 5: T cell distribution, differentiation and PD-1 expression on CD8$^+$ T cells in tumor-bearing NSG recipients after ACT of CD137$^+$ CD8$^+$ T cells.** A,B, frequency of CD3$^+$ T cells (% of human CD45$^+$ cells) at sacrifice in blood (**A**) or spleen (**B**) of tumor-bearing NSG mice after ACT of the indicated cell populations. C-E, CD8$^+$ T cell differentiation in blood (**C**), spleen (**D**) or TIL (**E**) of tumor-bearing NSG mice after ACT of indicated cell populations. **F**, frequency of PD-1 expression after *ex vivo* expansion of the indicated cell populations. G,H, frequency of PD-1 expression in blood (**G**) or spleen (**H**) of tumor-bearing NSG mice after ACT of indicated cell populations. Data are pooled from 1-4 independent experiments, n=3-16 mice per group. Significance by one-way ANOVA or 2way ANOVA, as appropriate. For each experiment, a different HPC donor was used for HIS mouse reconstitution and generation of autologous tumor. Data from individual experiments are indicated by different symbols, with individual mice from the same experiment indicated by the same symbol.
